# Consistent higher emissions of carbon dioxide, nitrous oxide, and methane during the daytime in reservoirs

**DOI:** 10.1101/2023.10.24.563772

**Authors:** E. León-Palmero, R. Morales-Baquero, I. Reche

## Abstract

Greenhouse gas (GHG) emissions from reservoirs are quantitatively relevant for atmospheric climatic forcing. These emissions have large temporal variability, with daily changes accounting for a substantial part of this variability. However, most estimations for GHG fluxes are based on the upscaling of discrete measurements performed during the daytime and, in general, they do not account for nighttime emissions. Here, we explored the daily patterns of CO_2_, N_2_O, and diffusive and ebullitive CH_4_ fluxes in two eutrophic reservoirs with contrasted morphometries in two different years. We found a daily pattern for CO_2_, N_2_O, and diffusive CH_4_ fluxes with consistent higher emissions during the daytime than during the nighttime, irrespectively of reservoir morphometry. These three diffusive fluxes showed evident daily synchrony suggesting a common driver. The emissions were coupled with the daily solar cycle, wind speed, and water temperature. The daily emissions of the CO_2_, N_2_O, and CH_4_ were also positive and significantly related to oxygen saturation. In contrast, we did not find a consistent daily pattern for the ebullitive CH_4_ fluxes, although they represented a significant fraction of the total CH_4_ emitted in these reservoirs. Our study suggests that the daily variability in GHG emissions may be as relevant as the variability at spatial scale or inter-system variability. Therefore, daily ranges should be considered in future GHG budgets to refine temporal trends of GHG emissions from reservoirs.

## 1. Introduction

The emission of greenhouse gases (GHG) such as CO_2_, CH_4_, and N_2_O from reservoirs is quantitatively relevant for atmospheric climatic forcing, with eutrophic reservoirs being responsible for most of this climatic forcing [1,2]. These emissions exhibit significant temporal (i.e., daily and seasonal) variability in reservoirs [3–6], with daily changes accounting for an important part of the total variability [3–5,7,8]. Daily patterns of GHG fluxes depend on the biological activity inside the reservoir, and on the physical and chemical processes that determine the CO_2_, CH_4_, and N_2_O concentrations and solubility, and their subsequent exchange with the atmosphere [4,9–12]. These environmental drivers can change considerably on a daily scale. The wind significantly affects water turbulence and promotes gas exchange across the air-water interface [4,12]. For this reason, gas transfer coefficients (k) are frequently parameterized as a function of the wind speed in emission estimates [13–15]. For instance, Liu *et al* [4] measured greater CO_2_ emissions in the nighttime due to the wind events at night. In contrast, the higher wind-induced turbulence during the daytime increased the emissions of diffusive and ebullitive CH_4_ in wetlands, reservoirs, and lakes [8,11,16,17]. Besides, temperature positively affects both GHG production (i.e., stimulating microbial metabolism) and emission (i.e., diminishing gas solubility). Daily changes in air and water temperature are frequently correlated to the daily pattern of diffusive emissions of CH_4_ [18] and N_2_O [10,19,20]. In addition, wind speed and cooling events can also promote upwelling motions and waterside convection events in shallow systems that can bring CO_2_ and CH_4_ from rich waters in deeper layers to the surface waters, increasing the emissions during the nighttime [21,22].

Solar radiation determines the daily cycle of photosynthesis and photochemistry in surface waters. Whereas photosynthesis (CO_2_ uptake) only occurs during the daytime, respiration (CO_2_ release) occurs throughout the whole day. The respiration of primary producers and heterotrophs may determine higher emissions of CO_2_ during the nighttime than during the daytime, depending on photosynthetic rates [4]. Moreover, CH_4_ emissions are closely linked to photosynthesis in lakes and reservoirs [1,23–27]. Bižić *et al* [27] demonstrated that the production of CH_4_ by cyanobacteria is associated with photosynthetic activity, following a daily cycle dependent on solar radiation. In relation to N_2_O production, the daily changes in the dissolved oxygen concentration affect the rates of microbial nitrogen processing during the nitrification and the coupling nitrification-denitrification [10,29,30], contributing to the diurnal patterns of N_2_O concentration and emission found in rivers [20,31,32]. On the other hand,, solar radiation also catalyzes photochemical reactions that decompose chromophoric and recalcitrant organic molecules into smaller ones, like carboxylic acids, that can be further completely mineralized up to CO_2_ and CH_4_ affecting their total production [34–40].

Most estimations for GHG fluxes are based on the upscaling of discrete measurements performed during the daytime and, usually, they did not account for the nighttime emissions [4,8]. Therefore, serious biases could affect the global temporal estimates of GHG emissions from reservoir. On this regard, some previous works have highlighted the diel variations in CO_2_ and CH_4_ fluxes [i.e., 3,8,41], but no study has evaluated yet these differences in N_2_O fluxes in lakes or reservoirs, where these emissions can be particularly high if the ecosystem receives large amounts of N from the watershed [2]. With the aim of contributing to fill this gap, here we described the daily patterns in CO_2_, N_2_O, and CH_4_ fluxes in two eutrophic reservoirs located in agricultural landscapes (i.e., with N loads especially high) with contrasting morphometric properties during the summer stratification in two different years. We discriminated the total CH_4_ emissions into the diffusive and ebullitive components. Finally, we determined the environmental drivers explaining these daily patterns, and analyzed its relevance in the context of temporal budgets.

## 2. Material and Methods

### 2.1. Study reservoirs

We performed three 24-hour sampling campaigns in two eutrophic reservoirs (Cubillas and Iznájar) located in southeastern Spain during the stratification period. Both reservoirs were built for irrigation and water supply purposes. We measured the GHG fluxes in Cubillas reservoir in 2016 (from July 14^th^ to July 15^th^) and in 2018 (from June 21^st^ to June 22^nd^), and in Iznájar reservoir in 2018 (July 8^th^ to July 9^th^). The Cubillas reservoir is a small and shallow system with a maximum capacity of 19 hm^3^, in which temporal variation in volume and maximum depth are related to annual rainfall. The year 2016 was dry (326.6 mm of rainfall and 7.0 meters of maximum depth) with an extreme summer, whereas the year 2018 was wet (509.4 mm of rainfall and 8.9 meters of maximum depth). The Iznájar reservoir is a bigger and deeper system than the Cubillas reservoir, with a maximum capacity of 981 hm^3^ and a mean depth of 22.4 m. We collected these data from The Confederación Hidrográfica del Guadalquivir (CHG; https://www.chguadalquivir.es/), and Infraestructura de Datos Espaciales de Andalucía (IDEAndalucia; http://www.ideandalucia.es/portal/web/ideandalucia/). We provided a further characterization of these two systems, and more information on GHG fluxes and concentrations measured in these reservoirs in previous studies [2,27,42].

### 2.2. Quantification of CO_2_, N_2_O, and CH_4_ fluxes

We measured CO_2_, CH_4_, and N_2_O fluxes from 13 to 24 times along the 24-hour sampling using a high-resolution laser-based Cavity Ring-Down Spectrometer (CRDS PICARRO G2508) coupled to a floating chamber [2]. We calculated the fluxes using the following equation (Eq. 1) [43]:

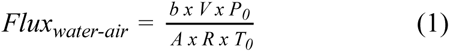

Where *Flux _water-air_* (μmol m^-2^ s^-1^) is the flux from the water surface to the atmosphere; the *b* (ppmv s^-1^) value is the slope of the linear regression between the time and the concentration of each gas inside the chamber; the *V* (m^3^) is the floating chamber volume; the *A* (m^2^) is the floating chamber area; the P_0_ (Pa) is the barometric pressure; the R is the gas constant (8.314 m^3^ Pa K^-1^ mol^-1^); and T_0_ (K) is the ambient temperature. We checked that the slope was significantly different from zero for each measurement using a two-tailed t-Student test. We also calculated the coefficient of determination (R^2^) for each measurement, accepting those whose R^2^ > 0.85 [2,44]. For CH_4_ fluxes, we computed the *b* value using the end-point concentrations and the time interval between them (Eq. 2) [43]:

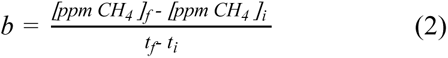

Where *[ppm CH_4_]_f_* and *[ppm CH_4_]_i_*are the CH_4_ concentration in the floating chamber at the end and the beginning of the time considered; and *t_f_* and *t_i_*are the time at the end and the beginning of the measurement, respectively. We discriminated the total CH_4_ fluxes (i.e., *CH_4_^total^*) in two components: the CH_4_ emitted by diffusion (*CH_4diffusion_*) and the CH_4_ emitted by ebullition (*CH_4ebullition_*). The *CH_4total_* was calculated following the eq 1, and the slope (b) was calculated following the eq 2. The CH_4_ flux discrimination between diffusion and ebullition in (*CH_4diffusion_*) and (*CH_4ebullition_*) components was performed using an algorithm inspired in the one proposed by Hoffmann *et al* [45], but adapting the procedure to higher data frequency. Briefly, the algorithm sets a variable moving window to generate several data subsets per each measurement, calculating the slope of a linear regression and different statistics for each subset. We established the moving window with a size of 50 data. The resulting slopes for each measurement are evaluated according to strict exclusion criteria to keep only the slopes for the diffusion component and filter out the ebullitive events. These inclusion criteria included: a significant regression slope between CH_4_ concentration and time (p < 0.05); determination coefficients R^2^ > 0.5; non-abrupt changes within each data subset (slope < 0.012, determined by data inspection); and the interquartile range test to discharge the slopes outside of the range between the upper and the lower quartile. Finally, we calculated the mean value for the resulting slopes that passed all the statistical tests. The resulting slope is used to calculate the diffusive component of the flux following the eq 1. The ebullitive component CH_4ebullition_ is estimated by subtracting the identified CH_4diffusion_ from the CH_4 total_ calculated before (eq. 3):

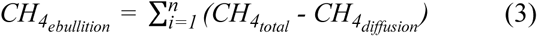

### 2.3. Physico-chemical characterization of water column and atmospheric conditions

We measured air temperature, barometric pressure (HANNA HI 9828), and wind speed (MASTECH MS6252A) at the beginning of each flux measurement. We recorded the dissolved oxygen concentration and saturation and the water temperature in the epilimnion every 10 minutes during the 24-hour sampling using a miniDOT (PME) submersible water logger. We also measured surface CH_4_, N_2_O, dissolved organic carbon (DOC), dissolved inorganic carbon (DIC), nitrate, nitrite, ammonia, and Chl-*a* concentrations once per day using standard methods. More details are provided in the Supporting Information (see Detailed Methods).

### 2.4. Circular statistics

Daytime is a circular variable measured at a closed and cyclical scale. Therefore, we used circular statistics, which is more appropriate to analyze variables with a cyclical nature [46]. We used the software Oriana to perform circular statistics and plot the data in rose diagrams [47]. We performed linear-circular correlation [48–50] between a circular variable (solar time) and a linear one (the fluxes or environmental variables). This correlation coefficient ranges from 0 to 1, so there is no negative correlation. We studied the synchrony between the greenhouse gas fluxes using the Spearman’s rank correlation analysis. We used R [51] for the rest of the statistical analysis and plots.

## 3. Results

### 3.1. Daily patterns in CO_2_, N_2_O, and CH_4_ fluxes

CO_2_ fluxes ranged from 0.10 to 0.42 µmol m^-2^ s^-1^ in Cubillas 2016, from 0.17 to 1.46 µmol m^-2^ s^-1^ in Cubillas 2018, and from -0.01 to 0.35 µmol m^-2^ s^-1^ in Iznájar. We measured the maximum values during the daytime, and the minimum values during the nighttime (Figure 1a-c, and Table S1). In 2018, we detected significant circular correlations between the solar time and the CO_2_ fluxes in the 24-hour cycles performed in Cubillas and Iznájar reservoirs (Figure 1b-c, and Table S2). N_2_O fluxes ranged more than one order of magnitude from 0.03 to 0.31 nmol m^-2^ s^-1^ in Cubillas 2016, from 0.08 to 1.13 nmol m^-2^ s^-1^ in Cubillas 2018, and from -0.09 to 0.46 nmol m^-2^ s^-1^ in Iznájar. Like in the CO_2_ emissions, we also measured the maximum values during the daytime, and the minimum values during the nighttime (Figure 1d-f, and Table S1). We detected significant circular correlations between the solar time and the N_2_O fluxes in the three 24-hour cycles performed (Figure 1d-f, and Table S2). The daily values of CH_4_ diffusive fluxes ranged from 0.00 to 989.15 nmol m^-2^ s^-1^ in Cubillas 2016, from 11.31 to 142.89 nmol m^-2^ s^-1^ in Cubillas 2018, and from 0.00 to 14.61 nmol m^-2^ s^-1^ in Iznájar (Figure 1g-i, and Table S1). In 2018, we also detected significant circular correlations between solar time and the diffusive CH_4_ emissions in Cubillas and Iznájar reservoirs (Figure 1h-i, and Table S2). For the ebullitive fraction, the values varied more than two orders of magnitude from 0.00 to 1815.32 nmol m^-2^ s^-1^ in Cubillas 2016, from 1.05 to 206.79 nmol m^-2^ s^-1^ in Cubillas 2018, and from 0.00 to 27.06 nmol m^-2^ s^-1^ in Iznájar (Figure 1j-l). We did not find any significant correlation between solar time and the ebullitive CH_4_ emissions (Table S2). The contribution of the ebullitive fraction to the total CH_4_ emission varied from 33 to 55 % during the daytime, and from 17 to 81 % during the nighttime (mean values, Table S1). Ebullitive fluxes contributed more to the total emissions in the shallower reservoir (Cubillas) than in the deeper one (Iznájar), reaching almost the 100 % of the total CH_4_ emission punctually in the Cubillas reservoir in 2016 (Table S1).

**Figure 1.**
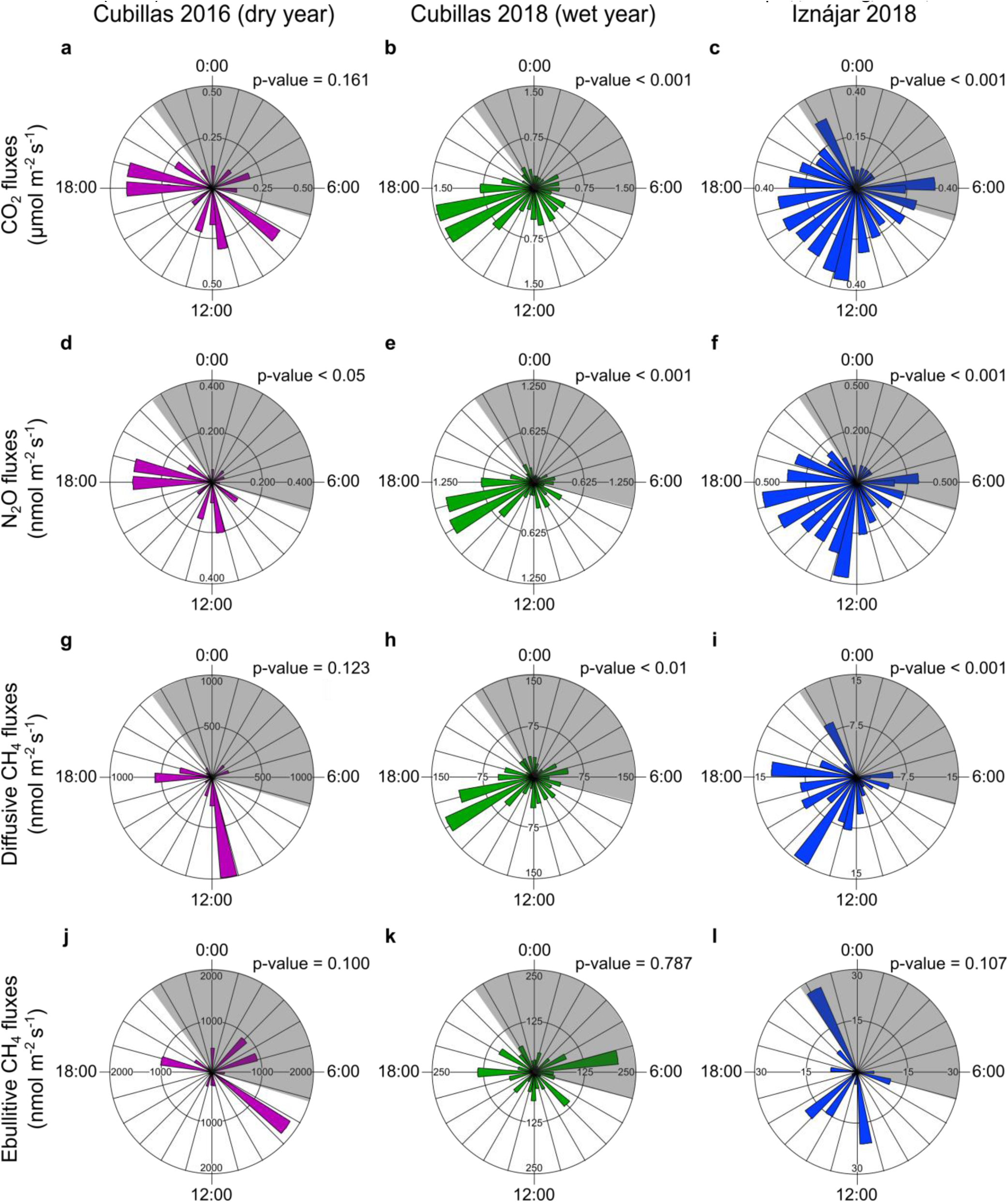
Rose diagrams of daily GHG fluxes in relation to solar time. We show the CO_2_ (a, b, c), N_2_O (d, e, f), diffusive CH_4_ (g, h, i), and ebullitive CH_4_ (j, k, l) fluxes in the Cubillas reservoir in 2016 (purple), in the Cubillas reservoir in 2018 (green), and in the Iznájar reservoir in 2018 (blue). Solar time is shown with numbers outside the circle, and numbers inside the circle show the GHG emission thresholds using different scales. The shaded areas refer to the night hours. We also provided the p-values for the circular-linear correlations between solar time (circular variable) and greenhouse gas fluxes (linear variable).

We found that the diffusive GHG fluxes were strongly synchronous in the three 24-hour measurement cycles performed, especially CO_2_ and N_2_O fluxes (Table S3). Besides, CO_2_ and N_2_O fluxes were higher in the Cubillas reservoir in 2018 than in 2016. In contrast, CH_4_ emissions were higher in Cubillas 2016 (i.e., the driest and warmest year) than in Cubillas 2018. Diffusive CH_4_ emissions were also significantly correlated to N_2_O fluxes in the three daily cycles and correlated to CO_2_ fluxes in the Cubillas and Iznájar reservoirs in 2018. In contrast, the ebullitive component of CH_4_ fluxes correlated to the CO_2_ fluxes only in the Iznájar reservoir in 2018.

### 3.2. Drivers of daily patterns of GHG fluxes

We studied wind speed (m s^-1^), water temperature (°C), and the percentage of oxygen saturation (%) (a biological indicator of the budget between photosynthesis and respiration in the surface waters) as environmental drivers for the GHG fluxes. We show their mean, minimum, and maximum values in Table S4. Based on circular-linear correlation analysis, the environmental drivers were significantly correlated to the solar time (Table S5). We show the daily cycle of wind speed, water temperature, oxygen saturation, and the GHG fluxes in Figure 2, and the results of the linear regressions between these environmental drivers and the GHG fluxes in Table 1.

**Figure 2.**
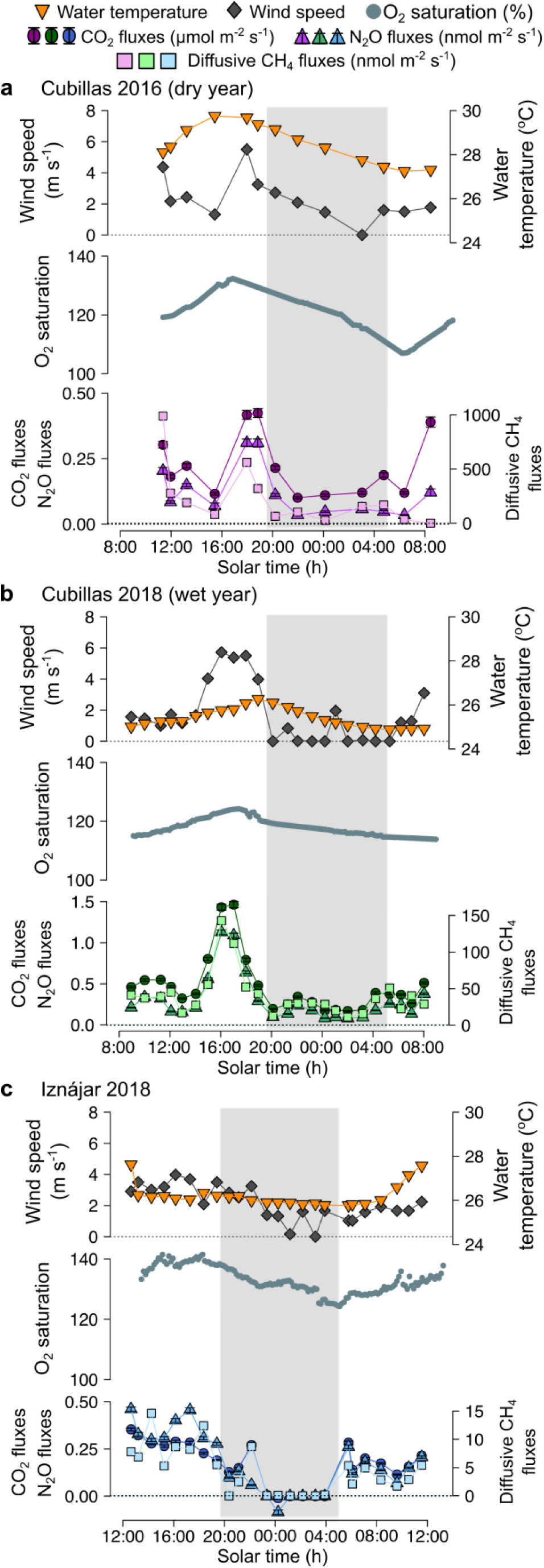
Wind speed, water temperature, and oxygen saturation as drivers for the variability of GHG fluxes at daily scales in Cubillas 2016 (a), Cubillas 2018 (b), and Iznájar 2018 (c). The shaded areas stand for the nighttime. Note the different scales.

**Table 1.** Results of the linear regressions between wind speed (m s^-1^), water temperature (°C), or oxygen saturation (%), and the GHG fluxes. CO_2_ emissions are provided in µmol m^-2^ s^-1^, and N_2_O and CH_4_ are provided in nmol m^-2^ s^-1^.

| GHG flux (y) | Driver (x) | n | Equation (estimate ± std. error) | Adj R<sup>2</sup> | p-value | Sampling |
| --- | --- | --- | --- | --- | --- | --- |
| $\text{CO}_2$ | Wind speed | 13 | $y = 0.08 (\pm 0.05) + 0.06 (\pm 0.02) x$ | 0.46 | < 0.01 | Cubillas 2016 |
| | | 24 | $y = 0.20 (\pm 0.05) + 0.16 (\pm 0.02) x$ | 0.70 | < 0.001 | Cubillas 2018 |
| | | 24 | $y = 0.00 (\pm 0.04) + 0.08 (\pm 0.02) x$ | 0.49 | < 0.001 | Iznájar 2018 |
|  | Water temperature | 13 | not significant |  | 0.548 | Cubillas 2016 |
| | | 24 | $y = -8.41 (\pm 2.52) + 0.35 (\pm 0.10) x$ | 0.33 | < 0.01 | Cubillas 2018 |
| | | 24 | $y = -1.79 (\pm 0.37) + 0.07 (\pm 0.01) x$ | 0.54 | < 0.001 | Iznájar 2018 |
|  | Oxygen saturation | 13 | not significant |  | 0.354 | Cubillas 2016 |
| | | 24 | $y = -8.90 (\pm 2.04) + 0.08 (\pm 0.02) x$ | 0.47 | < 0.001 | Cubillas 2018 |
| | | 24 | $y = -1.84 (\pm 0.66) + 0.02 (\pm 0.00) x$ | 0.26 | < 0.01 | Iznájar 2018 |
| $\text{N}_2\text{O}$ | Wind speed | 13 | $y = -0.01 (\pm 0.03) + 0.06 (\pm 0.01) x$ | 0.65 | < 0.001 | Cubillas 2016 |
| | | 24 | $y = 0.08 (\pm 0.04) + 0.13 (\pm 0.02) x$ | 0.71 | < 0.001 | Cubillas 2018 |
| | | 24 | $y = -0.04 (\pm 0.05) + 0.10 (\pm 0.02) x$ | 0.46 | < 0.001 | Iznájar 2018 |
|  | Water temperature | 13 | marginally significant |  | 0.082 | Cubillas 2016 |
| | | 24 | $y = -7.37 (\pm 2.02) + 0.30 (\pm 0.08) x$ | 0.37 | < 0.001 | Cubillas 2018 |
| | | 24 | $y = -2.59 (\pm 0.45) + 0.10 (\pm 0.02) x$ | 0.61 | < 0.001 | Iznájar 2018 |
| | Oxygen saturation | 13 | $y = -0.76 (\pm 0.40) + 0.01 (\pm 0.00) x$ | 0.25 | < 0.05 | Cubillas 2016 |
| | | 24 | $y = -7.45 (\pm 1.67) + 0.07 (\pm 0.01) x$ | 0.47 | < 0.001 | Cubillas 2018 |
| | | 24 | $y = -3.12 (\pm 0.78) + 0.02 (\pm 0.01) x$ | 0.42 | < 0.001 | Iznájar 2018 |
| Diffusive $\text{CH}_4$ | Wind speed | 13 | $y = -107.09 (\pm 105.10) + 144.65 (\pm 39.06) x$ | 0.51 | < 0.01 | Cubillas 2016 |
| | | 24 | $y = 20.18 (\pm 5.76) + 11.68 (\pm 2.27) x$ | 0.53 | < 0.001 | Cubillas 2018 |
| | | 24 | $y = -0.68 (\pm 1.53) + 2.37 (\pm 0.64) x$ | 0.36 | < 0.01 | Iznájar 2018 |
|  | Water temperature | 13 | not significant |  | 0.617 | Cubillas 2016 |
| | | 24 | $y = -702.92 (\pm 219.97) + 28.95 (\pm 8.57) x$ | 0.31 | < 0.01 | Cubillas 2018 |
| | | 24 | $y = -57.08 (\pm 14.78) + 2.31 (\pm 0.56) x$ | 0.42 | < 0.001 | Iznájar 2018 |
|  | Oxygen saturation | 13 | not significant |  | 0.478 | Cubillas 2016 |
| | | 24 | $y = -674.19 (\pm 192.96) + 6.07 (\pm 1.64) x$ | 0.36 | < 0.01 | Cubillas 2018 |
| | | 24 | $y = -76.74 (\pm 21.87) + 0.61 (\pm 0.16) x$ | 0.36 | < 0.01 | Iznájar 2018 |

In the Cubillas reservoir (2016 and 2018), wind speed was the main driver of the CO_2_, N_2_O, and diffusive CH_4_ fluxes, and the maximum emissions coincided with the maximum wind speeds (Figure 2, Table 1). However, in the Iznájar reservoir in 2018, water temperature showed a slightly better relationship to the CO_2_, N_2_O, and diffusive CH_4_ fluxes than wind speed (Figure 2, and Table 1). The variability range of wind speed in the Cubillas reservoir was higher than in the Iznájar reservoir, while water temperature showed higher variability in the Iznájar reservoir than in the Cubillas reservoir (Table S4). Consequently, the larger the variability range, the larger the explanatory power of a parameter is.

Oxygen saturation also showed a daily cycle presumably determined by the photosynthesis and respiration activity. The maximum values for oxygen saturation in the three daily cycles were closed to the maximum values for the emissions (Figure 2). We found that the oxygen saturation was positively related to the fluxes in most cases, with the more significant fits being for N_2_O fluxes (Table 1). We did not find any significant relationship with the ebullitive component of CH_4_ emission, only a marginally significant relationship to the water temperature in Cubillas 2016. In the linear regression models reported in Table 1, we detected that the slopes (estimate ± std. error) that explained the effect of wind speed, water temperature, and oxygen saturation were similar for the same 24-hour cycle for the CO_2_ and the N_2_O fluxes. In contrast, the slopes for the diffusive CH_4_ emissions were different from the CO_2_ and N_2_O (Table 1).

## 4. Discussion

### 4.1. Daily variability

Our data reveal that daily changes are a relevant component of the variability in the GHG fluxes in reservoirs. This significant variation along the day, especially in the CH_4_ fluxes, exceeded the inter-seasonal variation in the CO_2_, N_2_O, and CH_4_ fluxes previously reported in these two reservoirs between the summer stratification and the winter mixing [2]. Therefore, this daily variability must be considered when estimating temporal trends of GHG emissions and their climate forcing, and the reservoir carbon budgets between sedimentation and emissions. In this regard, we calculated how different time periods during a usual working day may overestimate or underestimate the fluxes of GHG in comparison to the 24 h measurements in these two reservoirs in 2018 (Table 2). These working hours corresponds to 9 am to 5 pm (equivalent to 7 am to 3 pm in solar time), in the time zone where the reservoirs are located (CEST, UTC + 02:00). We didn’t find a time range fitting the three study gases in the two reservoirs. Measuring GHG only from 9 to 5 pm may underestimate the fluxes by a 15% in Cubillas, and overestimate them by a 30% in Iznájar, because the maximum emissions were located at different times of the day. The period 9 – 2 pm, and 10 – 3 pm accurately represented daily CO_2_ and N_2_O mean emissions, respectively, in both reservoirs. Different time ranges from 10 to 5 pm closely aligned with the mean CH_4_ daily emissions in both reservoirs.

**Table 2.**
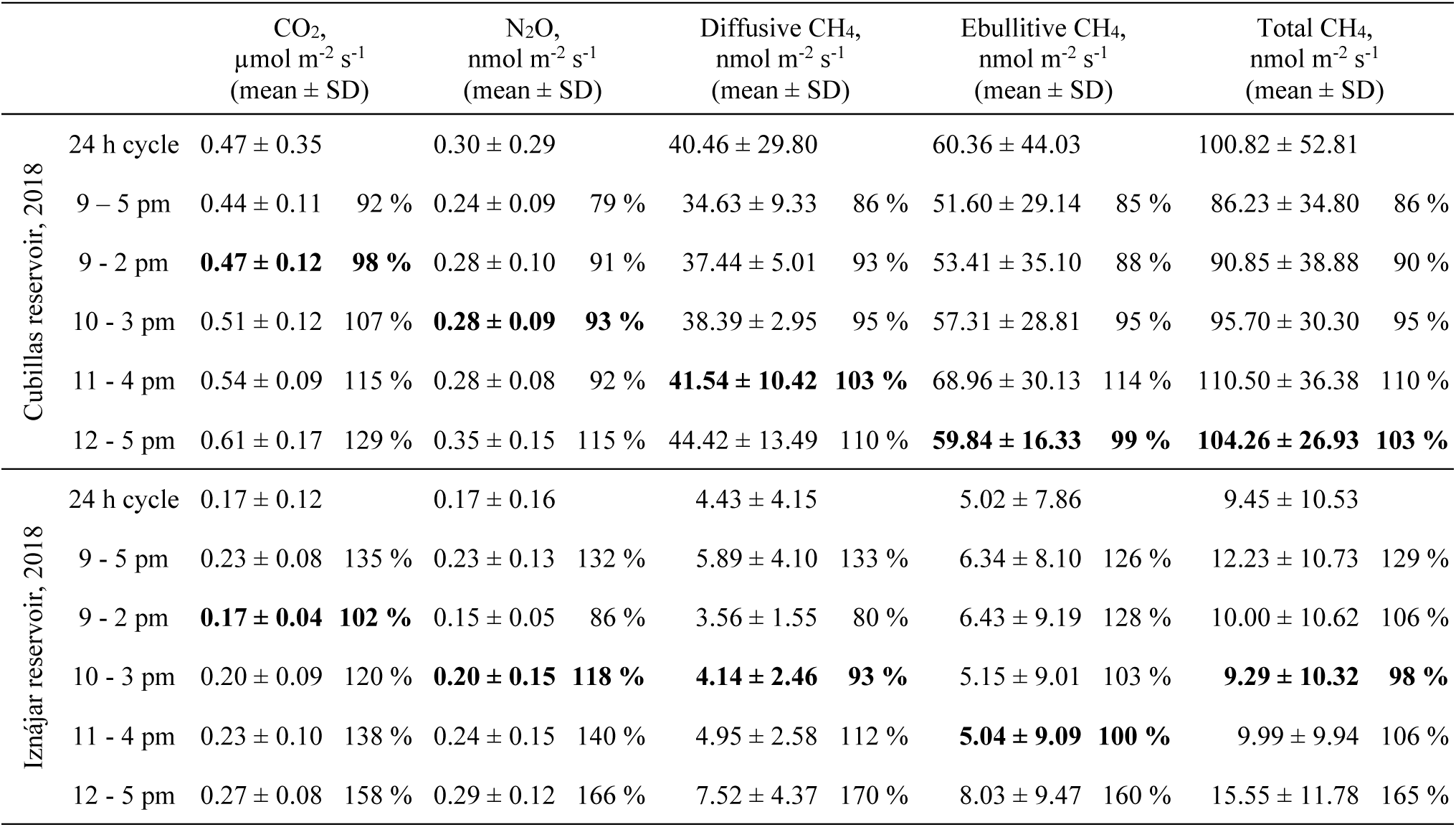
CO_2_, N_2_O, and CH_4_ fluxes during the 24 h cycles in 2018, and during specific daytime periods. The mean and standard deviation (SD) are provided. The percentage between the mean for that time ranges, and the 24 h mean is provided to evaluate the over or underestimation of that specific time period in relation to the daily mean. The best approximations are bolded.

|  |  | CO<sub>2</sub>,<br>μmol m<sup>-2</sup> s<sup>-1</sup><br>(mean ± SD) |  | N<sub>2</sub>O,<br>nmol m<sup>-2</sup> s<sup>-1</sup><br>(mean ± SD) |  | Diffusive CH<sub>4</sub>,<br>nmol m<sup>-2</sup> s<sup>-1</sup><br>(mean ± SD) |  | Ebullitive CH<sub>4</sub>,<br>nmol m<sup>-2</sup> s<sup>-1</sup><br>(mean ± SD) |  | Total CH<sub>4</sub>,<br>nmol m<sup>-2</sup> s<sup>-1</sup><br>(mean ± SD) |  |
| --- | --- | --- | --- | --- | --- | --- | --- | --- | --- | --- | --- |
| Cubillas reservoir, 2018 | 24 h cycle | 0.47 ± 0.35 |  | 0.30 ± 0.29 |  | 40.46 ± 29.80 |  | 60.36 ± 44.03 |  | 100.82 ± 52.81 |  |
|  | 9 – 5 pm | 0.44 ± 0.11 | 92 % | 0.24 ± 0.09 | 79 % | 34.63 ± 9.33 | 86 % | 51.60 ± 29.14 | 85 % | 86.23 ± 34.80 | 86 % |
|  | 9 – 2 pm | **0.47 ± 0.12** | **98 %** | 0.28 ± 0.10 | 91 % | 37.44 ± 5.01 | 93 % | 53.41 ± 35.10 | 88 % | 90.85 ± 38.88 | 90 % |
|  | 10 – 3 pm | 0.51 ± 0.12 | 107 % | **0.28 ± 0.09** | **93 %** | 38.39 ± 2.95 | 95 % | 57.31 ± 28.81 | 95 % | 95.70 ± 30.30 | 95 % |
|  | 11 – 4 pm | 0.54 ± 0.09 | 115 % | 0.28 ± 0.08 | 92 % | **41.54 ± 10.42** | **103 %** | 68.96 ± 30.13 | 114 % | 110.50 ± 36.38 | 110 % |
|  | 12 – 5 pm | 0.61 ± 0.17 | 129 % | 0.35 ± 0.15 | 115 % | 44.42 ± 13.49 | 110 % | **59.84 ± 16.33** | **99 %** | **104.26 ± 26.93** | **103 %** |
| Iznájar reservoir, 2018 | 24 h cycle | 0.17 ± 0.12 |  | 0.17 ± 0.16 |  | 4.43 ± 4.15 |  | 5.02 ± 7.86 |  | 9.45 ± 10.53 |  |
|  | 9 – 5 pm | 0.23 ± 0.08 | 135 % | 0.23 ± 0.13 | 132 % | 5.89 ± 4.10 | 133 % | 6.34 ± 8.10 | 126 % | 12.23 ± 10.73 | 129 % |
|  | 9 – 2 pm | **0.17 ± 0.04** | **102 %** | 0.15 ± 0.05 | 86 % | 3.56 ± 1.55 | 80 % | 6.43 ± 9.19 | 128 % | 10.00 ± 10.62 | 106 % |
|  | 10 – 3 pm | 0.20 ± 0.09 | 120 % | **0.20 ± 0.15** | **118 %** | **4.14 ± 2.46** | **93 %** | 5.15 ± 9.01 | 103 % | **9.29 ± 10.32** | **98 %** |
|  | 11 – 4 pm | 0.23 ± 0.10 | 138 % | 0.24 ± 0.15 | 140 % | 4.95 ± 2.58 | 112 % | **5.04 ± 9.09** | **100 %** | 9.99 ± 9.94 | 106 % |
|  | 12 – 5 pm | 0.27 ± 0.08 | 158 % | 0.29 ± 0.12 | 166 % | 7.52 ± 4.37 | 170 % | 8.03 ± 9.47 | 160 % | 15.55 ± 11.78 | 165 % |

We summarized the studies on the daily variability of the GHG emissions in reservoirs and lakes in Table S6, including some investigations on rivers, wetlands, and soils. Previous studies found similar ranges of daily CO_2_ fluxes in deep reservoirs and lakes [3,41], but a higher variability in systems shallower than the study reservoirs [52,53]. In rivers and soils, the daily variability of the N_2_O fluxes was up to one order of magnitude [19,32,54,55]. No previous studies on the daily variability of N_2_O fluxes in lakes or reservoirs were found. Concerning the variability of the CH_4_ emissions, it was up to one order of magnitude in reservoirs and lakes [3,11,41,56], but higher in shallow systems [22,53,57].

### 4.2. Drivers of the CO_2_, N_2_O and CH_4_ fluxes

Wind speed was the main driver of CO_2_, N_2_O, and diffusive CH_4_ daily fluxes in Cubillas, while water temperature was the main driver in Iznájar. We detected a higher range of wind speed in Cubillas (2016 and 2018) than in Iznájar and a higher range in the water temperature in Iznájar than in Cubillas. These differences in the variation range of the environmental drivers may explain the differences between the corresponding drivers of the GHG fluxes in these two reservoirs. The wind speed influenced CO_2_ and N_2_O emissions with similar intensity, as suggested by the fact that the slopes of these relationships were similar (Table 1). However, the slope for the diffusive fluxes of CH_4_ was variable between years and reservoirs (Table 1).

The higher CO_2_ emissions during the daytime than during the nighttime appear initially counterintuitive, considering that photosynthesis occurs during the daytime. Previous works found larger CO_2_ emissions in the nighttime in shallow systems disturbed by high winds and convective processes [4,52,53], but higher CO_2_ emissions during the daytime than during the nighttime in deep systems [41,58,59] (Table S6). In the study reservoirs the CO_2_ emissions were determined by the lithology of the watershed (i.e., most dissolved inorganic carbon derived from carbonate rocks weathering), instead of the photosynthesis-respiration cycle [2]. On the other hand, solar radiation determines the direct CO_2_ photoproduction and indirect promotion of bacterial mineralization throughout photobleaching, that can increase the surface concentration of CO_2_ and could explain, to some extent, the higher CO_2_ emissions in daytime [34–39,60].

The N_2_O emissions were also higher during the daytime than during the nighttime. Previous studies also reported higher N_2_O emissions in the daytime in rivers [19,20,30–32,55], and in soils [54,61], but we did not find studies on N_2_O fluxes at daily scales in lakes or reservoirs (Table S6). In the study reservoirs, the daily pattern in the N_2_O emissions was correlated to wind speed, water temperature, and oxygen saturation, similarly to rivers [19,20,55]. Previous studies stated that the daily changes in the dissolved oxygen can affect microbial nitrogen processing rates during nitrification and coupled nitrification-denitrification [10,29,30].

We also detected higher emissions of diffusive CH_4_ during the daytime than during the nighttime, which is consistent with previous studies in stratified and mixed lakes [8,59,62], and in shallower systems [11,57]. In contrast, Martinez-Cruz *et al* [41] did not detect differences in the CH_4_ emissions at daily scales (Table S6). The higher wind speed during the daytime can cause more effective turbulent transfer and higher emissions of diffusive CH_4_ than during the nighttime in lakes and wetlands [8,11,16,59,63]. However, some shallow systems can be also affected by convective mixing during the nighttime, showing the opposite pattern [21,22]. Microbial CH_4_ production can be also affected by the daily cycle. Methanogenesis is strongly temperature-dependent [64], which may contribute to explaining the positive correlation between daily changes in water temperature and the diffusive CH_4_ emissions [18]. Besides, solar radiation can also enhance aerobic CH_4_ production in surface waters [28] and inhibit CH_4_ oxidation [65]. Therefore, higher CH_4_ production and lower CH_4_ oxidation in surface waters exposed to solar radiation may result in larger CH_4_ emissions at daytime.

We did not find a well-defined daily pattern for CH_4_ ebullitive emissions, which were not correlated to the other GHG emissions, or the environmental variables considered in this study. In contrast, other studies measured higher ebullitive fluxes during the daytime, due to the higher winds, which promoted water movements and CH_4_ bubbling [17,18]. Ebullitive emissions can represent a relevant fraction of total CH_4_ emissions [11,17,18]. For this reason, to determine the environmental factors that control these ebullition events is challenging and needs further exploration.

The daily patterns of the CO_2_, N_2_O, and diffusive CH_4_ fluxes were recurrent, despite the differences in magnitude between years and reservoirs. These differences in the magnitude of the fluxes in the Cubillas reservoir in 2016 and 2018 are similar to the differences detected in the DIC, N_2_O, and CH_4_ concentration in the surface waters (Table S4). In a previous study in 12 reservoirs, including Cubillas and Iznájar, we detected that CO_2_ and N_2_O fluxes depended on the carbonates and nitrogen inputs from their watersheds, respectively [2]. The year 2018 was wetter than 2016, with a higher runoff, and inputs of DIC and nitrogen into the reservoirs. In fact, we detected higher concentrations of DIC, nitrate, and dissolved N_2_O in 2018 than in 2016 (Table S4), that may determine the higher fluxes of CO_2_, and N_2_O [2]. In contrast, the CH_4_ fluxes were an order of magnitude higher in 2016 than in 2018, reflecting the differences in the CH_4_ concentration in surface waters (Table S4). In 2016, Cubillas had warmer and shallower waters, and a higher Chl-*a* concentration than in 2018 (Table S4), and that could explain the higher concentration and emission of CH_4_ in the year 2016 than in 2018. Previous studies showed that the concentration and emission of CH_4_ are related to Chl-*a* concentration, mean depth of the system, and water temperature [2,17,27,66–68]. In addition, lower mean depth in 2016 than in 2018 also promoted CH_4_ ebullition [17,69,70].

### 4.3. Daily synchrony of fluxes and environmental drivers

The three diffusive fluxes showed evident daily synchrony, with higher fluxes during the daytime than during the nighttime, and they were coupled to the wind speed, water temperature, and oxygen saturation. This daily synchrony indicates the existence of an extrinsic force (i.e., acting from outside the system) driving the variation in the GHG emissions at daily scales. Initially, the environmental drivers were significantly related to solar time (Table S5). This correlation was transferred to the GHG fluxes themselves and caused that they were also significantly related to the solar time (and to the environmental drivers). We detected that CO_2_, N_2_O, and diffusive CH_4_ fluxes were significantly related to the solar time, except for CO_2_ and diffusive CH_4_ fluxes in the Cubillas reservoir in 2016. The lower number of measurements in 2016 (n = 13) than in 2018 (n = 24) could have caused the absence of statistically significant results during this year (Table S2). The fluxes of N_2_O were always significantly related to solar time. In addition to the three environmental drivers analyzed, solar radiation can also promote the abiotic and biotic production of greenhouse gases, as explained above. That can occur by promoting the direct and indirect photoproduction of CO_2_ [34–39,60], or by driving the photosynthesis – respiration cycle that also affects CH_4_ production (in addition to CO_2_) [28].

## Conclusions

We detected higher emissions of CO_2_, N_2_O, and diffusive CH_4_ during the daytime than during the nighttime. This daily pattern of the GHG fluxes was recurrent, despite the differences in magnitude between years and reservoirs. However, ebullitive fluxes, which are a relevant fraction of total CH_4_ emissions, did not show a clear daily pattern. The significant variability in the GHG emissions at daily scales demonstrates that daily differences should be considered when addressing temporal trends of GHG fluxes and their climatic forcing, and the carbon budgets between emissions and sedimentation in reservoirs. In addition, the synchrony of the GHG emissions at daily scales may be a common feature occurring in aquatic systems, but it has not been detected before because we still miss studies addressing simultaneously the fluxes of CO_2_, N_2_O, and CH_4_. In fact, this study has been the first one showing the significant differences in N_2_O emissions at daily scales in lentic systems.

## Author Contributions

The manuscript was written through contributions of all authors. ELP, RM-B, and IR contributed to data acquisition during the reservoir samplings, and EL-P processed the data. EL-P, RM-B and IR analyzed the data and discussed the results. EL-P wrote the first draft manuscript, which was complemented by significant contributions of RM-B and IR. IR and RM-B designed the study and obtained the funds. All authors have given approval to the final version of the manuscript.

## Acknowledgements

We especially thank Lucía Franco, Alba Contreras-Ruiz, Ángel Marcos Vicente, and Eulogio Corral for helping in the field, Jesús Forja and Teodora Ortega for helping with gas chromatography analysis at the Department of Physical Chemistry of the University of Cádiz. We thank the Hydrological Confederations of Guadalquivir and the Agencia Andaluza del Medio Ambiente y Agua (AMAYA) for facilitating the reservoir sampling. This research has been supported by the Spanish Ministry of Science, Innovation and Universities (grant no. RTI2018-098849-B-I00); the Consejería de Universidad, Investigación e Innovación from Junta de Andalucía and the European Regional Development Fund (ERDF; grant no. B-RNM-558-UGR20). Elizabeth León-Palmero was supported by a PhD fellowship from the Spanish Ministry of Education, Culture and Sport (grant nos. FPU014/02917), and later by a Marie Skłodowska-Curie postdoctoral fellowship (HORIZON-MSCA-2021-PF-01, project number: 101066750) by the European Commission at Princeton University.

## Data Availability Statement

Data supporting the findings of this study are available within article, and in the Supporting Information. Raw data are available on request from the authors.

## Supporting Information content

### Detailed methods

#### N_2_O, and CH_4_ concentrations in surface waters

We collected surface water (0.5 m) using a 5-l UWITEC sampling bottle for the chemical and biological analysis explained below. Samples for the dissolved N_2_O and CH_4_ analysis were carefully collected in air-tight Winkler bottles by duplicate once per day, preserved with a solution of HgCl_2_ (final concentration 1mM) to inhibit biological activity and sealed with Apiezon® grease to prevent gas exchange. We stored the samples in the dark until analysis in the laboratory. We measured dissolved N_2_O and CH_4_ using headspace equilibration in a 50 ml air-tight glass syringe (Agilent P/N 5190–1547) in duplicate from each bottle [27,42]. We took a quantity of 25 g of water (± 0.01 g) using the air-tight syringe and added a quantity of 25 mL of a standard gas mixture that has a methane and nitrous oxide concentration similar to atmospheric values (0.3 and 1.8 ppmv, respectively) to complete the volume of the syringe. The syringes were shaken for 5 min (VIBROMATIC Selecta) to ensure a complete mixing, and we waited for 5 min to reach the equilibrium. Finally, the gas in the syringe (≈ 20 mL) was injected manually in the gas chromatograph (Bruker® GC-450). The gas chromatograph was equipped with hydrogen flame ionization detector and electron capture detector to measure the concentration of N_2_O and CH_4_ simultaneously. We daily calibrated the detectors using three standard gas mixtures with N_2_O concentrations of 305, 474, 2000 ppbv, and CH_4_ concentrations of 1952, 10064, and 103829 ppbv, made and certified by Air Liquide (France). We calculated the gas concentration in the water samples from the concentration measured in the headspace using the Bunsen functions for the N_2_O [71], and for the CH_4_ solubility [72].

#### Dissolved C, N, and Chl-*a* analysis in the water column

Water samples for chemical and biological analysis were maintained at 4 °C until arrival at the laboratory. Then, we filtered 500 to 2000 mL of water through 0.7 *μ*m pore-size Whatman GF/F glass-fiber filters. We kept the filter to determine Chl-*a* concentration, and the filtered water for dissolved nutrients analysis. We acidified with phosphoric acid (final pH < 2) the samples for DOC. We measured DOC, and dissolved inorganic carbon (DIC) by high–temperature catalytic oxidation using a Shimadzu total organic carbon (TOC) analyzer (Model TOC-V CSH) [73]. The instrument was calibrated using a four-point standard curve of dried potassium hydrogen phthalate for DOC, and dried sodium bicarbonate and sodium carbonate for DIC. We analyzed two replicates and three to five injections per replicate for each sample. Samples for DOC analysis were purged with phosphoric acid for 20 min to eliminate DIC. We measured the NO_3_^−^ concentration using the ultraviolet spectrophotometric method, using a Perkin Elmer UV-Lambda 40 spectrophotometer at wavelengths of 220 nm and correcting for DOC absorbance at 275 nm [74]. We also determined the concentrations of NH ^+^and NO ^−^ by inductively coupled plasma–optical emission spectrometry (ICP-OES). We determined Chl-*a* concentration using the previous glass-fiber filters. We extracted the pigments from the filters with 95% methanol in the dark at 4C for 24 h [74]. We measured Chl-*a* absorption at the wavelength of 665 nm using a Perkin Elmer UV-Lambda 40 spectrophotometer, and we corrected the solution scattering at 750 nm.

### Supporting Tables

**Table S1.**
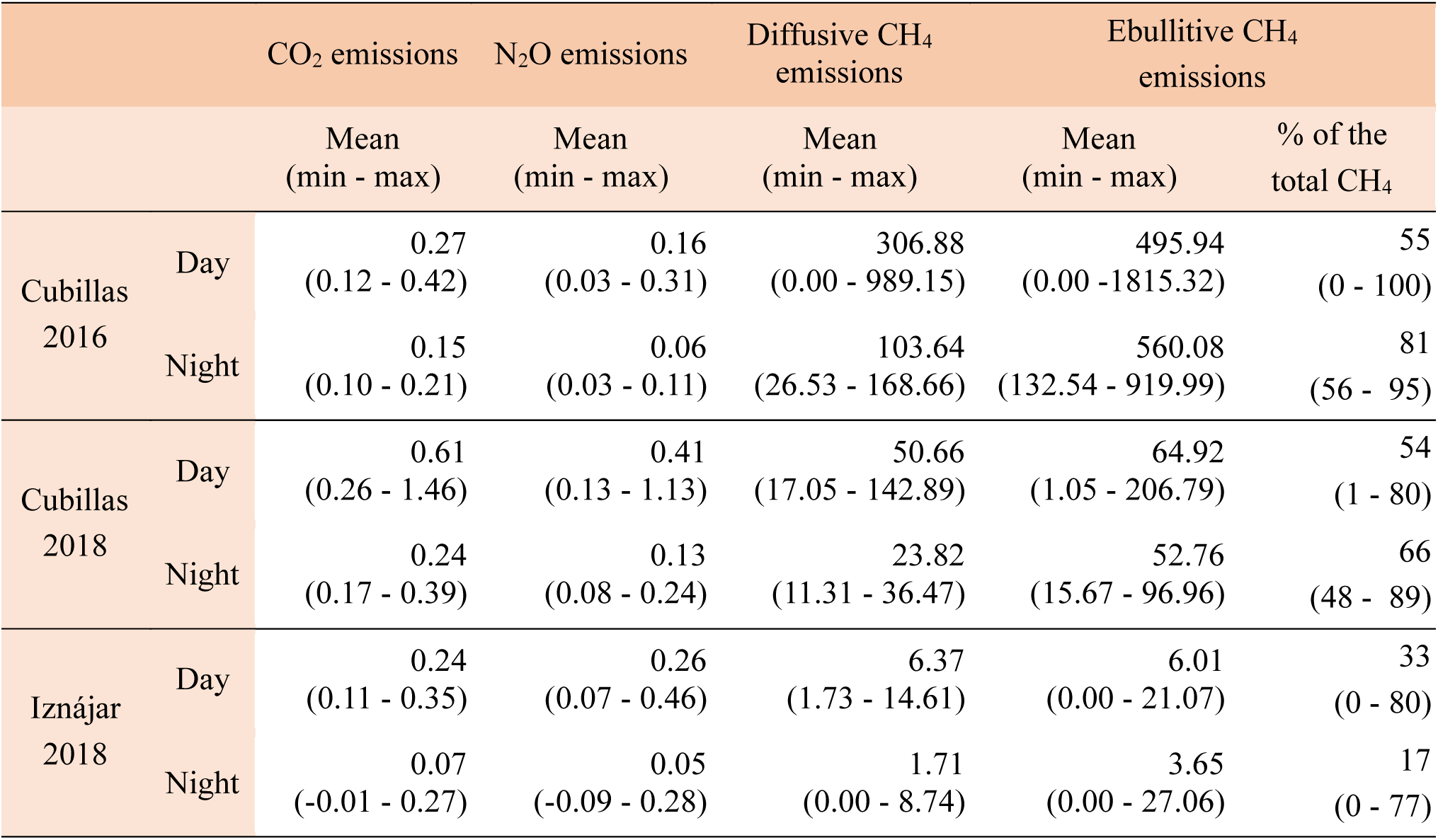
Emissions of CO_2_ (µmol m^-2^ s^-1^), N_2_O (nmol m^-2^ s^-1^), and CH_4_ by diffusion and ebullition (nmol m^-2^ s^-1^).

**Table S2.**
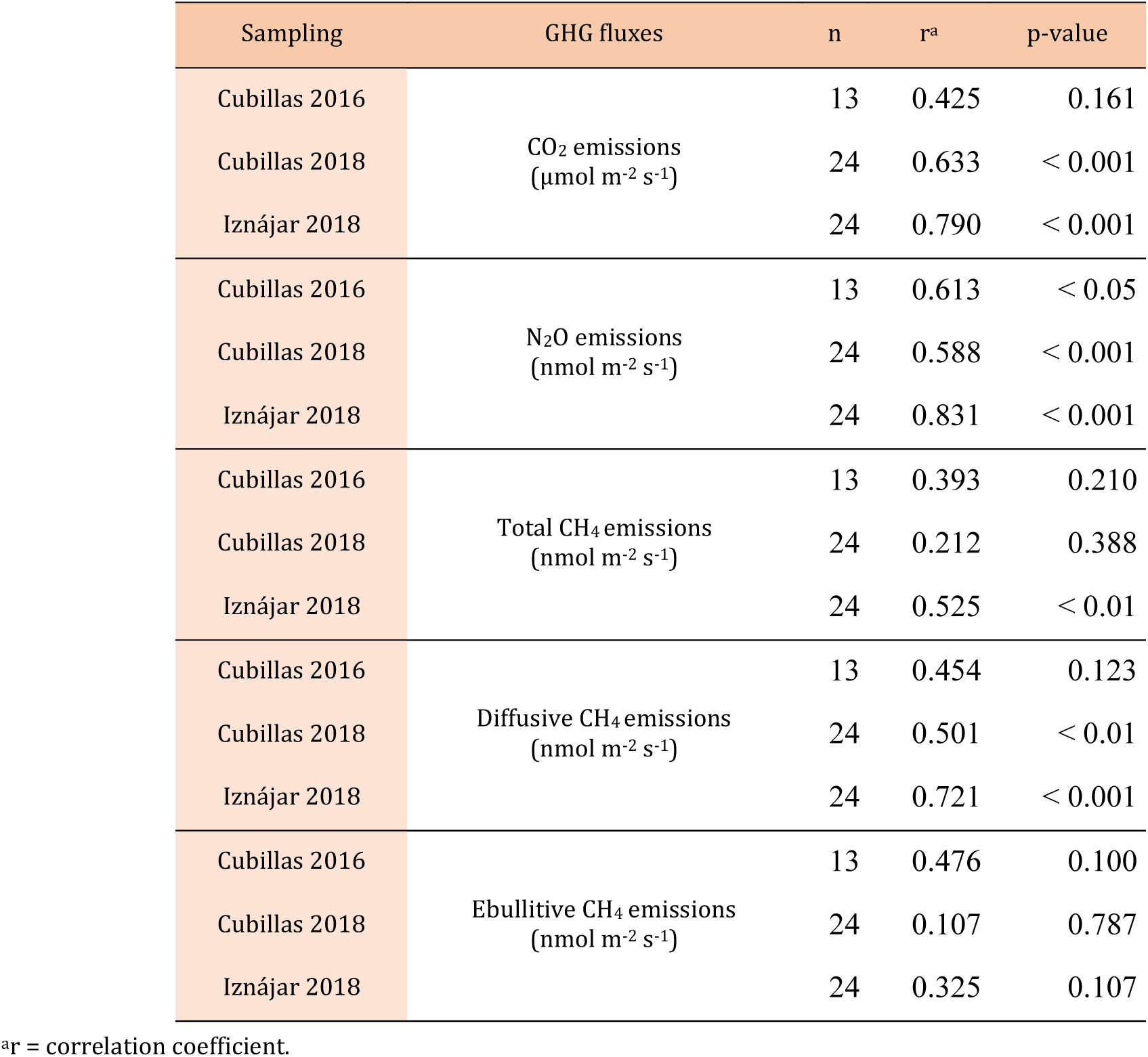
Statistical details of the circular-linear correlation between solar time (h, circular variable) and greenhouse gas fluxes (linear variable).

**Table S3.**
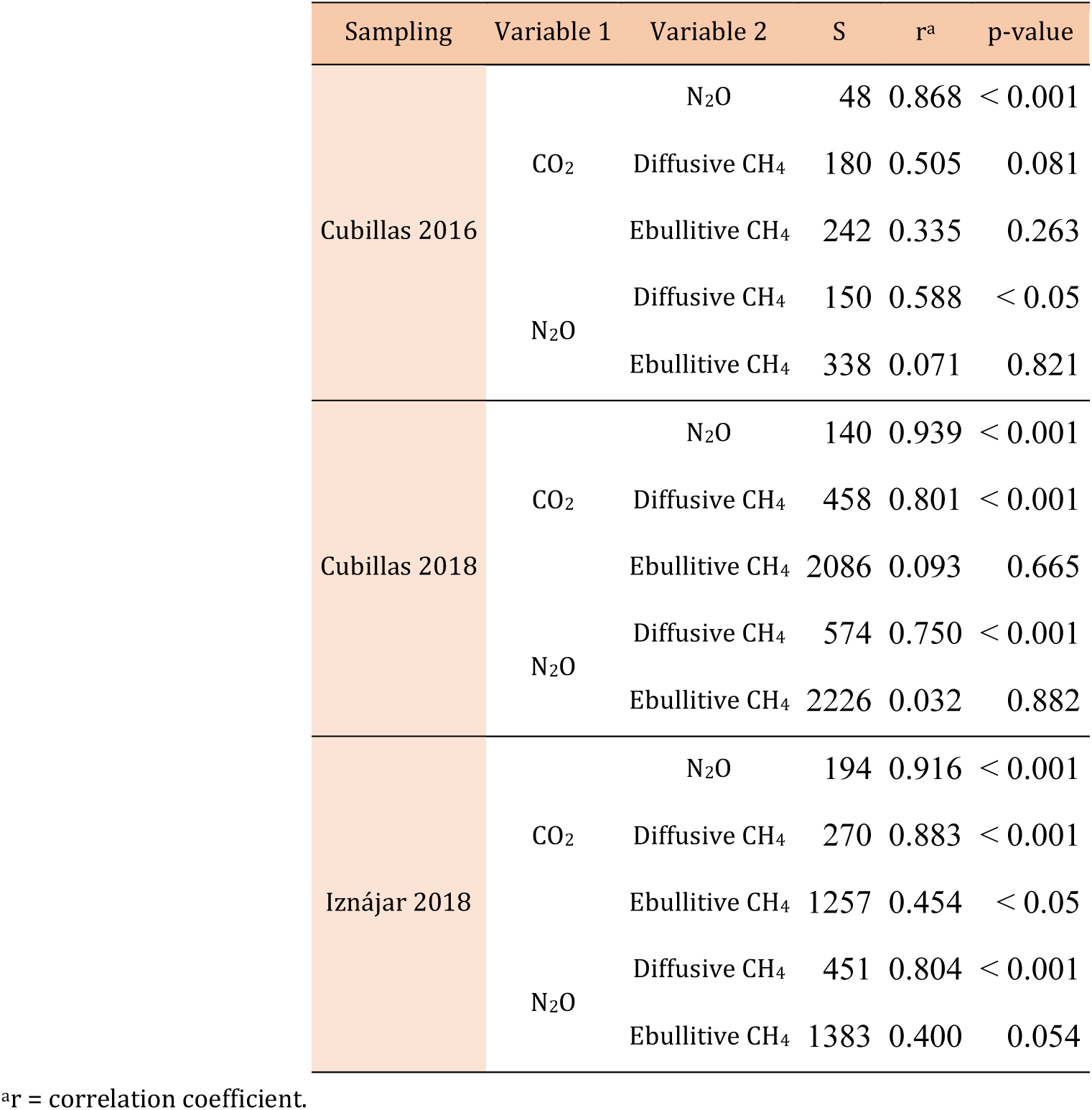
Statistical details of the Spearman’s correlation between greenhouse gas fluxes. CO_2_ emissions are provided in µmol m^-2^ s^-1^, and N_2_O and CH_4_ are provided in nmol m^-2^ s^-1^.

**Table S4.**
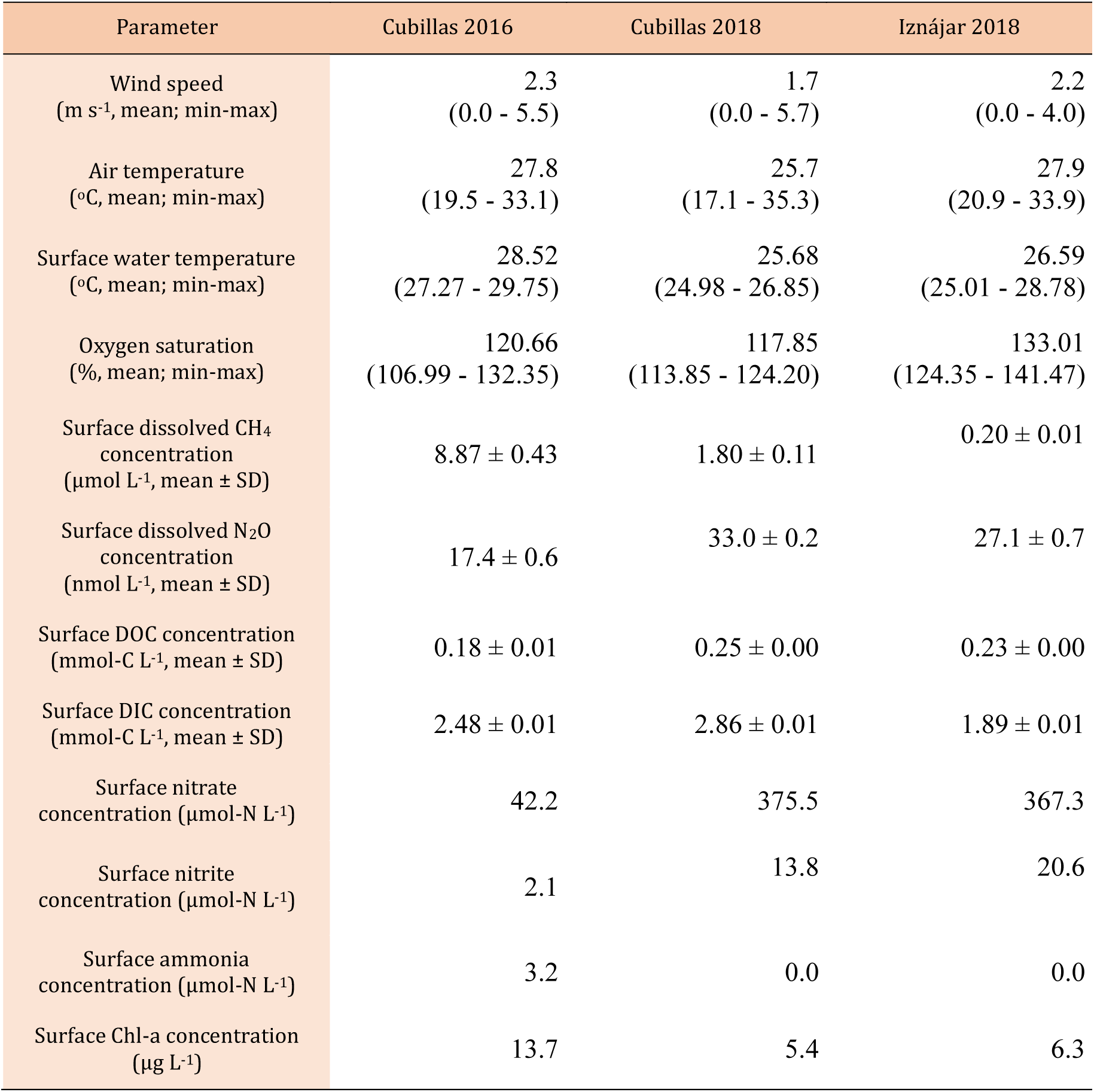
Physical, chemical, and biological parameters in Cubillas and Iznájar reservoirs. We provide the mean, minimum, and maximum values for wind speed, air temperature, surface water temperature, the oxygen saturation measured during the 24-hour sampling campaigns in Cubillas reservoir in July 2016 and June 2018, and in Iznájar reservoir in July 2018. For the rest of parameters, we provide discrete measurements for surface waters. n.m. = not measured.

**Table S5.**
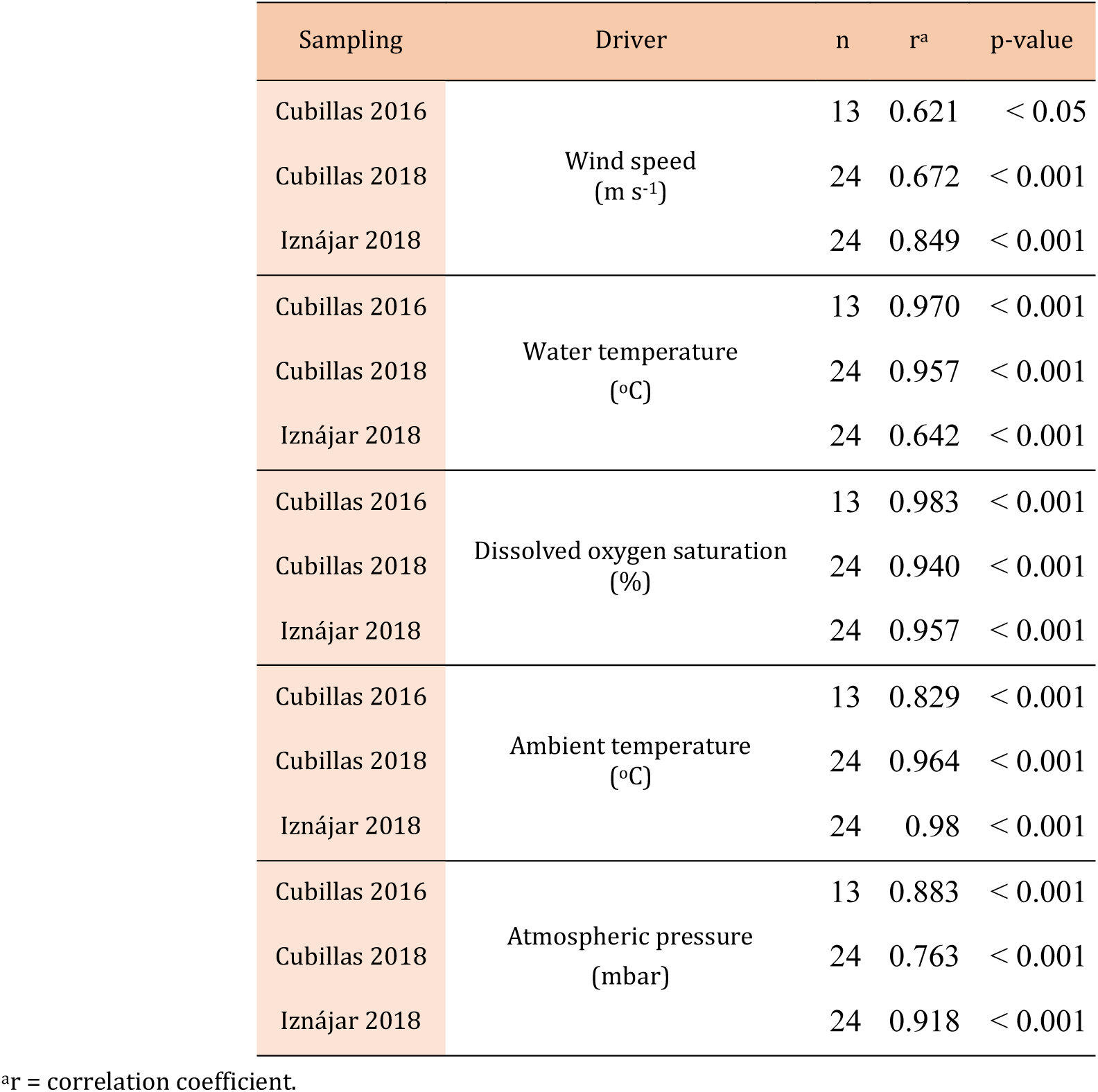
Statistical details of the circular-linear correlation between solar time (h, circular variable) and environmental drivers (linear variable).

**Table S6.**
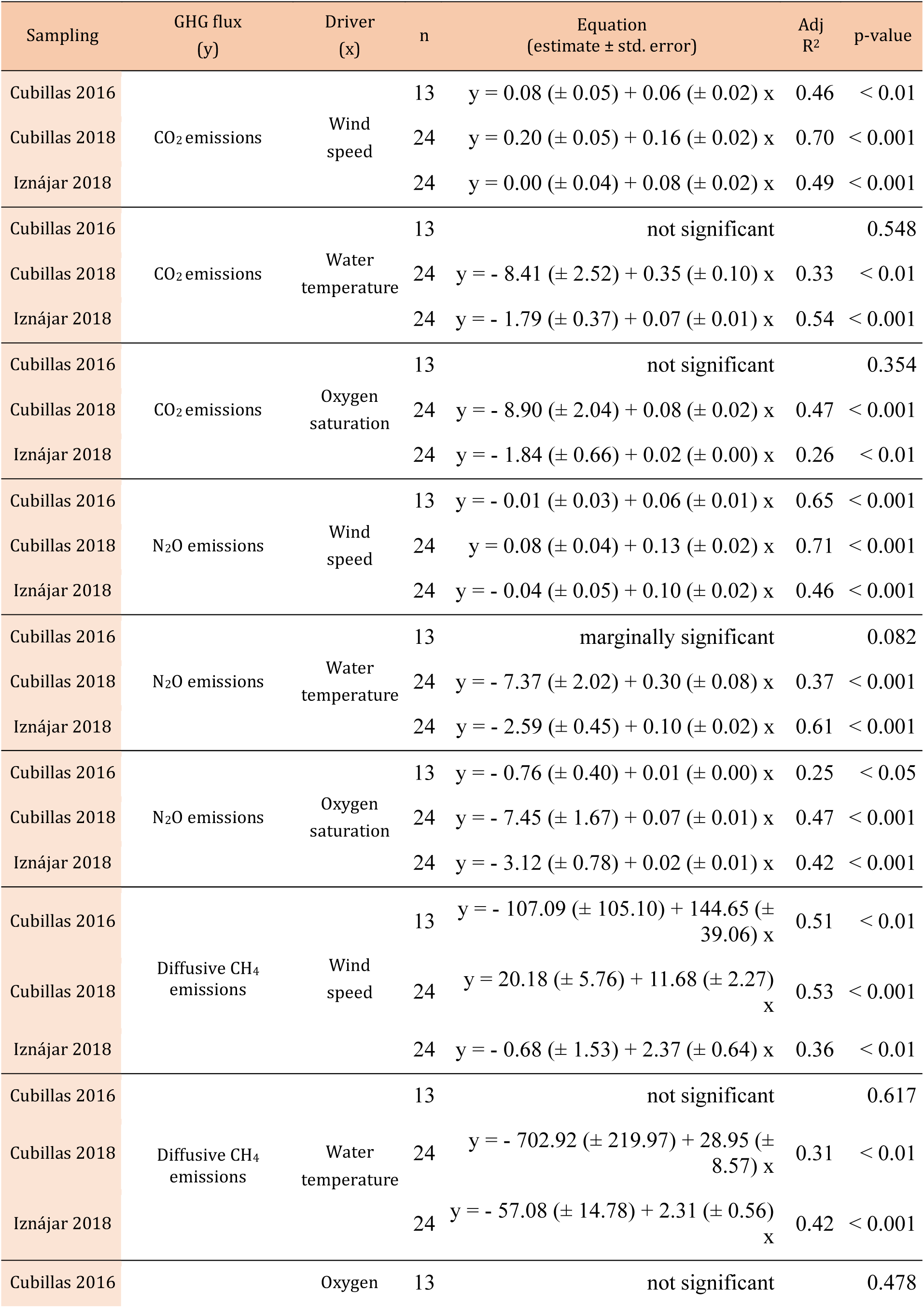
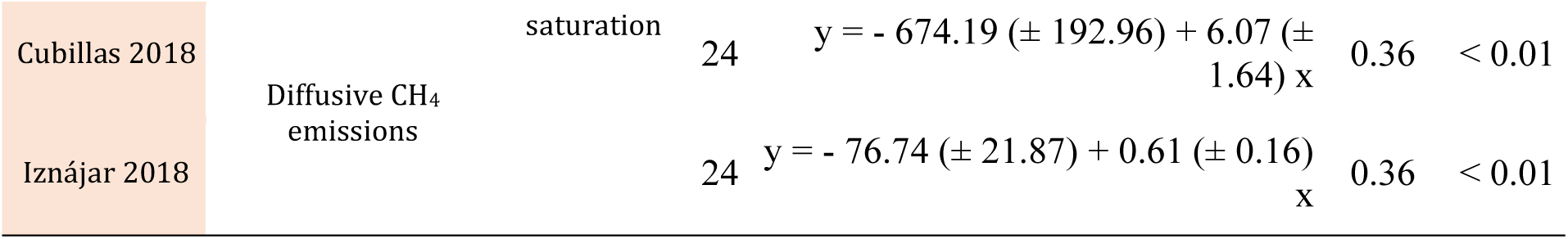
Results of the linear regressions between wind speed (m s^-1^), water temperature (°C), or oxygen saturation (%), and the GHG fluxes. CO_2_ emissions are provided in µmol m^-2^ s^-1^, and N_2_O and CH_4_ are provided in nmol m^-2^ s^-1^.

**Table S7.**
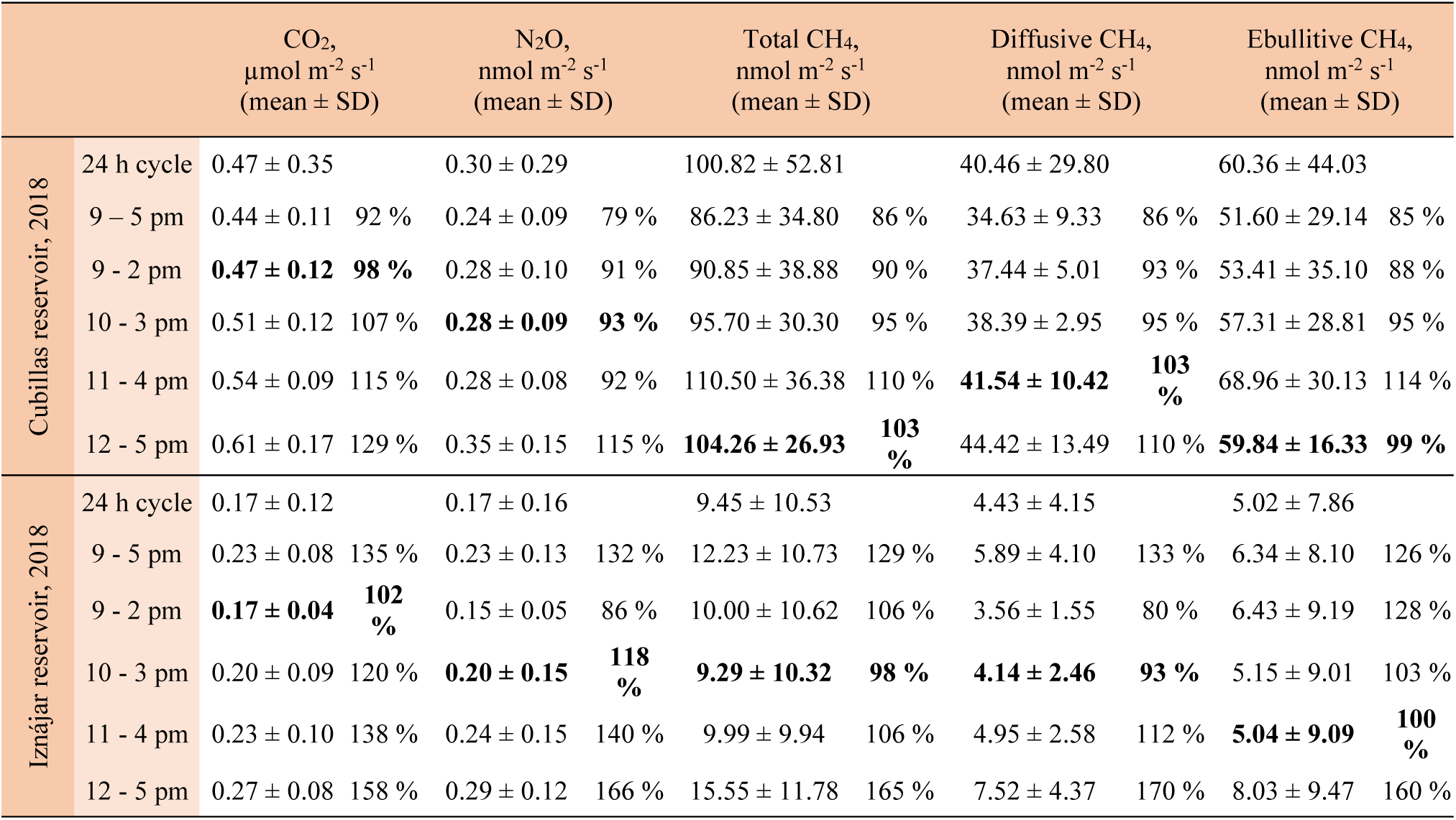
The fluxes of CO_2_, N_2_O, and CH_4_ during the 24 h cycles in 2018, and during specific daytime periods. The mean and standard deviation (SD) are provided. The percentage between the mean for that time period, and the 24 h mean is provided in order to evaluate the over or underestimation of that specific time period in relation to the daily mean.

**Table S8.**
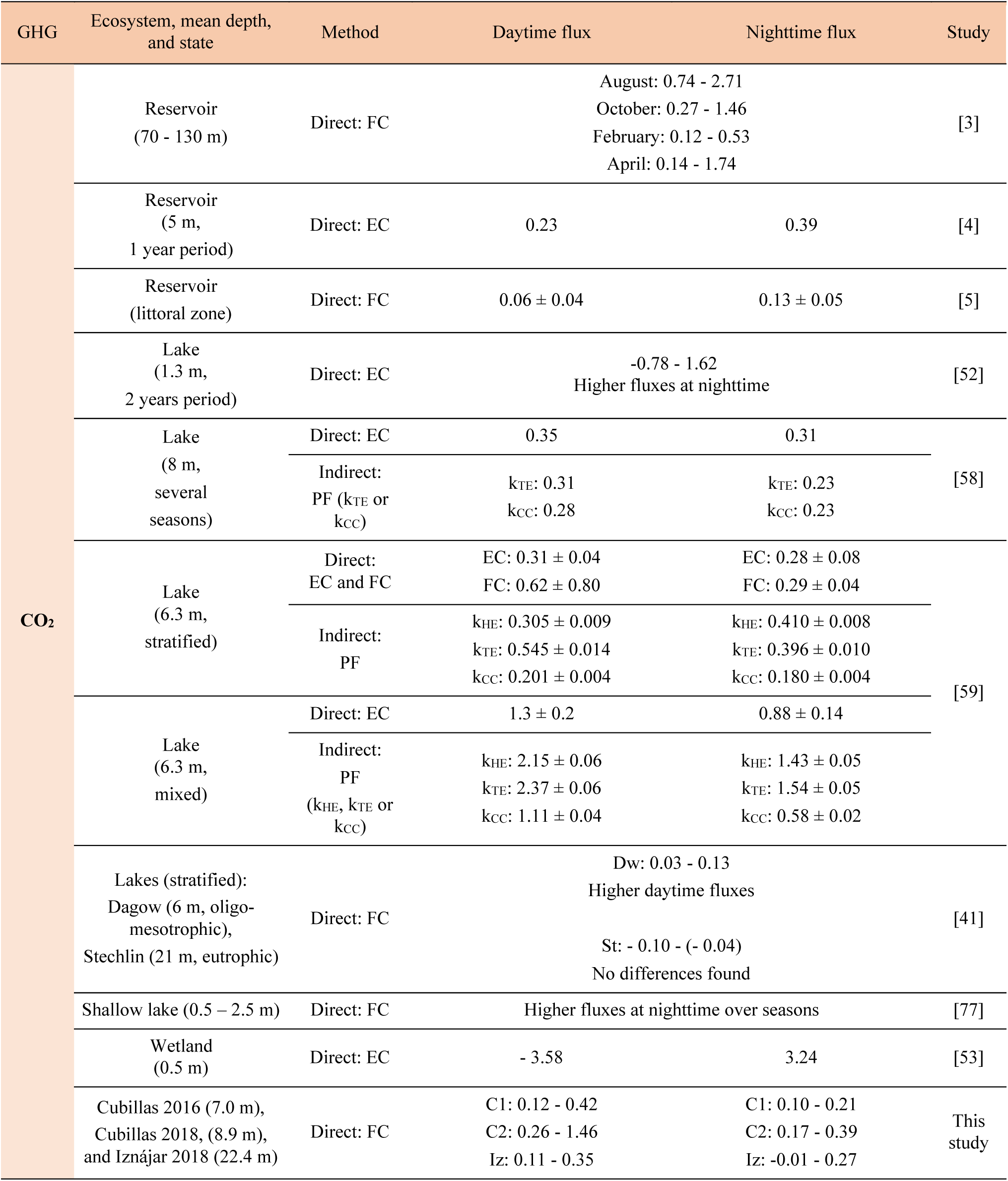
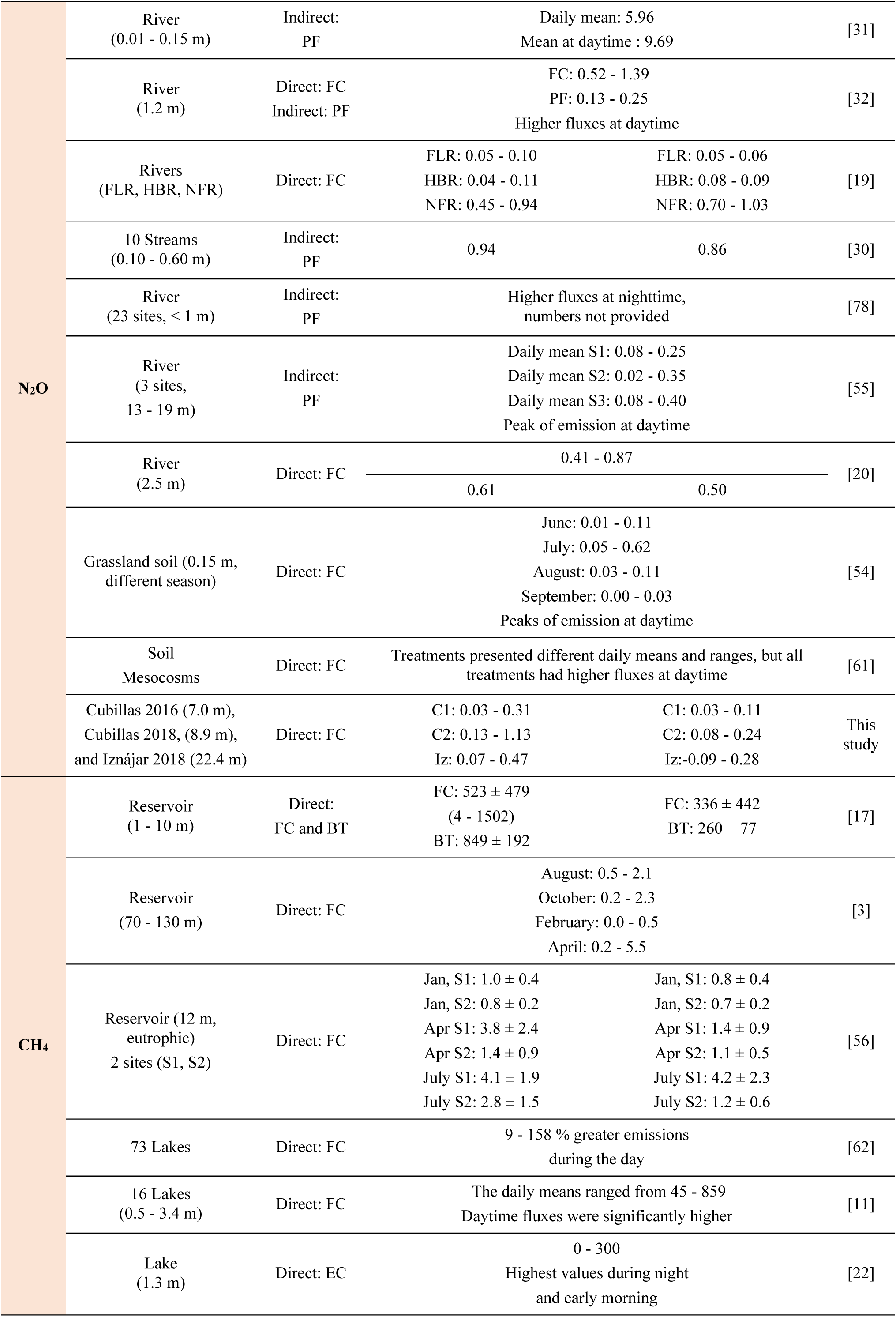
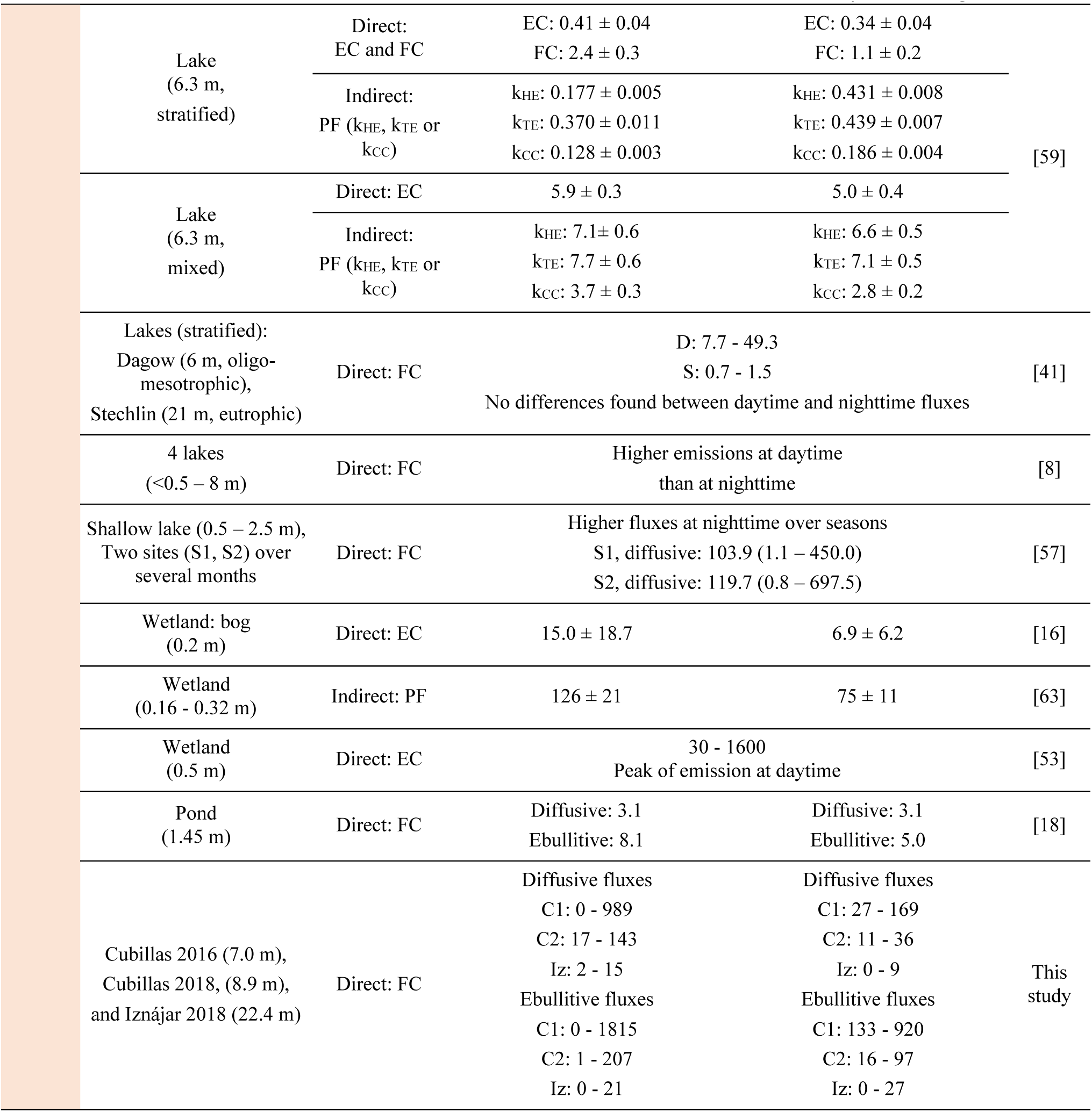
Summary table of published studies on the variability of CO_2_ (µmol m^-2^ s^-1^), N_2_O (nmol m^-2^ s^-1^), and CH_4_ (nmol m^-2^ s^-1^) fluxes at daily scales in different ecosystems and using different methods. We classified these methods in direct measurement, by eddy covariance technique (EC); closed/floating chamber (FC) and bubble traps for ebullitive fluxes (BT); and the indirect method that predicted the fluxes (PF) based on gas transfer velocities (k). The transfer velocities are traditionally based on wind-speed, as k_CC_ [13] and k_CW_ for low wind speeds [14]. Other gas transfer velocities also include the effects of water-side cooling to the gas transfer besides shear-induced turbulence, as k_HE_ [75] and k_TE_ [76]. Other authors preferred an empirical model for k, derived from laboratory experiments. We provided the daytime and nighttime mean (value ± standard error); or the range of variation (minimum - maximum). We provided the daily values when the data for daytime/nighttime are not supplied in the reference study.

