## Supporting Information for "Consistent higher emissions of carbon dioxide, nitrous oxide, and methane during the daytime in reservoirs"

Journal **XX** (XXXX) XXXXXX https://doi.org/XXXX/XXXX

Received xxxxxx
Accepted for publication xxxxxx
Published xxxxxx

### Supporting Information content:

Number of pages: 20
Detailed methods
Tables S1 – S8

 © xxxx IOP Publishing Ltd

 © xxxx IOP Publishing Ltd

Journal **XX** (XXXX) XXXXXX https://doi.org/XXXX/XXXX

### Supporting Tables

**Table S1.** Emissions of $CO_2$ (µmol m$^{-2}$ s$^{-1}$), $N_2O$ (nmol m$^{-2}$ s$^{-1}$), and $CH_4$ by diffusion and ebullition (nmol m$^{-2}$ s$^{-1}$).

| | | $CO_2$ emissions | $N_2O$ emissions | Diffusive $CH_4$ emissions | Ebullitive $CH_4$ emissions | |
|---|---|---|---|---|---|---|
| | | Mean (min - max) | Mean (min - max) | Mean (min - max) | Mean (min - max) | % of the total $CH_4$ |
| Cubillas 2016 | Day | 0.27 (0.12 - 0.42) | 0.16 (0.03 - 0.31) | 306.88 (0.00 - 989.15) | 495.94 (0.00 -1815.32) | 55 (0 - 100) |
| | Night | 0.15 (0.10 - 0.21) | 0.06 (0.03 - 0.11) | 103.64 (26.53 - 168.66) | 560.08 (132.54 - 919.99) | 81 (56 - 95) |
| Cubillas 2018 | Day | 0.61 (0.26 - 1.46) | 0.41 (0.13 - 1.13) | 50.66 (17.05 - 142.89) | 64.92 (1.05 - 206.79) | 54 (1 - 80) |
| | Night | 0.24 (0.17 - 0.39) | 0.13 (0.08 - 0.24) | 23.82 (11.31 - 36.47) | 52.76 (15.67 - 96.96) | 66 (48 - 89) |
| Iznájar 2018 | Day | 0.24 (0.11 - 0.35) | 0.26 (0.07 - 0.46) | 6.37 (1.73 - 14.61) | 6.01 (0.00 - 21.07) | 33 (0 - 80) |
| | Night | 0.07 (-0.01 - 0.27) | 0.05 (-0.09 - 0.28) | 1.71 (0.00 - 8.74) | 3.65 (0.00 - 27.06) | 17 (0 - 77) |


© xxxx IOP Publishing Ltd

**Table S2.** Statistical details of the circular-linear correlation between solar time (h, circular variable) and greenhouse gas fluxes (linear variable).

| Sampling | GHG fluxes | n | r[a] | p-value |
|---|---|---|---|---|
| Cubillas 2016 | $CO_2$ emissions (μmol $m^{-2}$ $s^{-1}$) | 13 | 0.425 | 0.161 |
| Cubillas 2018 | | 24 | 0.633 | < 0.001 |
| Iznájar 2018 | | 24 | 0.790 | < 0.001 |
| Cubillas 2016 | $N_2O$ emissions (nmol $m^{-2}$ $s^{-1}$) | 13 | 0.613 | < 0.05 |
| Cubillas 2018 | | 24 | 0.588 | < 0.001 |
| Iznájar 2018 | | 24 | 0.831 | < 0.001 |
| Cubillas 2016 | Total $CH_4$ emissions (nmol $m^{-2}$ $s^{-1}$) | 13 | 0.393 | 0.210 |
| Cubillas 2018 | | 24 | 0.212 | 0.388 |
| Iznájar 2018 | | 24 | 0.525 | < 0.01 |
| Cubillas 2016 | Diffusive $CH_4$ emissions (nmol $m^{-2}$ $s^{-1}$) | 13 | 0.454 | 0.123 |
| Cubillas 2018 | | 24 | 0.501 | < 0.01 |
| Iznájar 2018 | | 24 | 0.721 | < 0.001 |
| Cubillas 2016 | Ebullitive $CH_4$ emissions (nmol $m^{-2}$ $s^{-1}$) | 13 | 0.476 | 0.100 |
| Cubillas 2018 | | 24 | 0.107 | 0.787 |
| Iznájar 2018 | | 24 | 0.325 | 0.107 |

[a]r = correlation coefficient.

 © xxxx IOP Publishing Ltd

**Table S3.** Statistical details of the Spearman's correlation between greenhouse gas fluxes. $CO_2$ emissions are provided in µmol $m^{-2}$ $s^{-1}$, and $N_2O$ and $CH_4$ are provided in nmol $m^{-2}$ $s^{-1}$.

| Sampling | Variable 1 | Variable 2 | S | r[a] | p-value |
|---|---|---|---|---|---|
| Cubillas 2016 | $CO_2$ | $N_2O$ | 48 | 0.868 | < 0.001 |
| | | Diffusive $CH_4$ | 180 | 0.505 | 0.081 |
| | | Ebullitive $CH_4$ | 242 | 0.335 | 0.263 |
| | $N_2O$ | Diffusive $CH_4$ | 150 | 0.588 | < 0.05 |
| | | Ebullitive $CH_4$ | 338 | 0.071 | 0.821 |
| Cubillas 2018 | $CO_2$ | $N_2O$ | 140 | 0.939 | < 0.001 |
| | | Diffusive $CH_4$ | 458 | 0.801 | < 0.001 |
| | | Ebullitive $CH_4$ | 2086 | 0.093 | 0.665 |
| | $N_2O$ | Diffusive $CH_4$ | 574 | 0.750 | < 0.001 |
| | | Ebullitive $CH_4$ | 2226 | 0.032 | 0.882 |
| Iznájar 2018 | $CO_2$ | $N_2O$ | 194 | 0.916 | < 0.001 |
| | | Diffusive $CH_4$ | 270 | 0.883 | < 0.001 |
| | | Ebullitive $CH_4$ | 1257 | 0.454 | < 0.05 |
| | $N_2O$ | Diffusive $CH_4$ | 451 | 0.804 | < 0.001 |
| | | Ebullitive $CH_4$ | 1383 | 0.400 | 0.054 |

[a]r = correlation coefficient.

 © xxxx IOP Publishing Ltd

**Table S4.** Physical, chemical, and biological parameters in Cubillas and Iznájar reservoirs. We provide the mean, minimum, and maximum values for wind speed, air temperature, surface water temperature, the oxygen saturation measured during the 24-hour sampling campaigns in Cubillas reservoir in July 2016 and June 2018, and in Iznájar reservoir in July 2018. For the rest of parameters, we provide discrete measurements for surface waters. n.m. = not measured.

| Parameter | Cubillas 2016 | Cubillas 2018 | Iznájar 2018 |
|---|---|---|---|
| Wind speed ($m\ s^{-1}$, mean; min-max) | 2.3 (0.0 - 5.5) | 1.7 (0.0 - 5.7) | 2.2 (0.0 - 4.0) |
| Air temperature ($^{o}C$, mean; min-max) | 27.8 (19.5 - 33.1) | 25.7 (17.1 - 35.3) | 27.9 (20.9 - 33.9) |
| Surface water temperature ($^{o}C$, mean; min-max) | 28.52 (27.27 - 29.75) | 25.68 (24.98 - 26.85) | 26.59 (25.01 - 28.78) |
| Oxygen saturation (%, mean; min-max) | 120.66 (106.99 - 132.35) | 117.85 (113.85 - 124.20) | 133.01 (124.35 - 141.47) |
| Surface dissolved $CH_4$ concentration ($\mu mol\ L^{-1}$, mean ± SD) | 8.87 ± 0.43 | 1.80 ± 0.11 | 0.20 ± 0.01 |
| Surface dissolved $N_2O$ concentration ($nmol\ L^{-1}$, mean ± SD) | 17.4 ± 0.6 | 33.0 ± 0.2 | 27.1 ± 0.7 |
| Surface DOC concentration ($mmol\text{-}C\ L^{-1}$, mean ± SD) | 0.18 ± 0.01 | 0.25 ± 0.00 | 0.23 ± 0.00 |
| Surface DIC concentration ($mmol\text{-}C\ L^{-1}$, mean ± SD) | 2.48 ± 0.01 | 2.86 ± 0.01 | 1.89 ± 0.01 |
| Surface nitrate concentration ($\mu mol\text{-}N\ L^{-1}$) | 42.2 | 375.5 | 367.3 |
| Surface nitrite concentration ($\mu mol\text{-}N\ L^{-1}$) | 2.1 | 13.8 | 20.6 |
| Surface ammonia concentration ($\mu mol\text{-}N\ L^{-1}$) | 3.2 | 0.0 | 0.0 |
| Surface Chl-a concentration ($\mu g\ L^{-1}$) | 13.7 | 5.4 | 6.3 |


© xxxx IOP Publishing Ltd

Journal **XX** (XXXX) XXXXXX https://doi.org/XXXX/XXXX

**Table S5.** Statistical details of the circular-linear correlation between solar time (h, circular variable) and environmental drivers (linear variable).

| Sampling | Driver | n | r[a] | p-value |
|---|---|---|---|---|
| Cubillas 2016 | Wind speed ($m\ s^{-1}$) | 13 | 0.621 | < 0.05 |
| Cubillas 2018 | | 24 | 0.672 | < 0.001 |
| Iznájar 2018 | | 24 | 0.849 | < 0.001 |
| Cubillas 2016 | Water temperature (ºC) | 13 | 0.970 | < 0.001 |
| Cubillas 2018 | | 24 | 0.957 | < 0.001 |
| Iznájar 2018 | | 24 | 0.642 | < 0.001 |
| Cubillas 2016 | Dissolved oxygen saturation (%) | 13 | 0.983 | < 0.001 |
| Cubillas 2018 | | 24 | 0.940 | < 0.001 |
| Iznájar 2018 | | 24 | 0.957 | < 0.001 |
| Cubillas 2016 | Ambient temperature (ºC) | 13 | 0.829 | < 0.001 |
| Cubillas 2018 | | 24 | 0.964 | < 0.001 |
| Iznájar 2018 | | 24 | 0.98 | < 0.001 |
| Cubillas 2016 | Atmospheric pressure (mbar) | 13 | 0.883 | < 0.001 |
| Cubillas 2018 | | 24 | 0.763 | < 0.001 |
| Iznájar 2018 | | 24 | 0.918 | < 0.001 |

[a]r = correlation coefficient.


© xxxx IOP Publishing Ltd

Journal **XX** (XXXX) XXXXXX https://doi.org/XXXX/XXXX

**Table S6.** Results of the linear regressions between wind speed (m $s^{-1}$), water temperature (ºC), or oxygen saturation (%), and the GHG fluxes. $CO_2$ emissions are provided in µmol $m^{-2}$ $s^{-1}$, and $N_2O$ and $CH_4$ are provided in nmol $m^{-2}$ $s^{-1}$.

| Sampling | GHG flux (y) | Driver (x) | n | Equation (estimate ± std. error) | Adj $R^2$ | p-value |
|---|---|---|---|---|---|---|
| Cubillas 2016 | | | 13 | y = 0.08 (± 0.05) + 0.06 (± 0.02) x | 0.46 | < 0.01 |
| Cubillas 2018 | $CO_2$ emissions | Wind speed | 24 | y = 0.20 (± 0.05) + 0.16 (± 0.02) x | 0.70 | < 0.001 |
| Iznájar 2018 | | | 24 | y = 0.00 (± 0.04) + 0.08 (± 0.02) x | 0.49 | < 0.001 |
| Cubillas 2016 | | | 13 | not significant | | 0.548 |
| Cubillas 2018 | $CO_2$ emissions | Water temperature | 24 | y = - 8.41 (± 2.52) + 0.35 (± 0.10) x | 0.33 | < 0.01 |
| Iznájar 2018 | | | 24 | y = - 1.79 (± 0.37) + 0.07 (± 0.01) x | 0.54 | < 0.001 |
| Cubillas 2016 | | | 13 | not significant | | 0.354 |
| Cubillas 2018 | $CO_2$ emissions | Oxygen saturation | 24 | y = - 8.90 (± 2.04) + 0.08 (± 0.02) x | 0.47 | < 0.001 |
| Iznájar 2018 | | | 24 | y = - 1.84 (± 0.66) + 0.02 (± 0.00) x | 0.26 | < 0.01 |
| Cubillas 2016 | | | 13 | y = - 0.01 (± 0.03) + 0.06 (± 0.01) x | 0.65 | < 0.001 |
| Cubillas 2018 | $N_2O$ emissions | Wind speed | 24 | y = 0.08 (± 0.04) + 0.13 (± 0.02) x | 0.71 | < 0.001 |
| Iznájar 2018 | | | 24 | y = - 0.04 (± 0.05) + 0.10 (± 0.02) x | 0.46 | < 0.001 |
| Cubillas 2016 | | | 13 | marginally significant | | 0.082 |
| Cubillas 2018 | $N_2O$ emissions | Water temperature | 24 | y = - 7.37 (± 2.02) + 0.30 (± 0.08) x | 0.37 | < 0.001 |
| Iznájar 2018 | | | 24 | y = - 2.59 (± 0.45) + 0.10 (± 0.02) x | 0.61 | < 0.001 |
| Cubillas 2016 | | | 13 | y = - 0.76 (± 0.40) + 0.01 (± 0.00) x | 0.25 | < 0.05 |
| Cubillas 2018 | $N_2O$ emissions | Oxygen saturation | 24 | y = - 7.45 (± 1.67) + 0.07 (± 0.01) x | 0.47 | < 0.001 |
| Iznájar 2018 | | | 24 | y = - 3.12 (± 0.78) + 0.02 (± 0.01) x | 0.42 | < 0.001 |
| Cubillas 2016 | | | 13 | y = - 107.09 (± 105.10) + 144.65 (± 39.06) x | 0.51 | < 0.01 |
| Cubillas 2018 | Diffusive $CH_4$ emissions | Wind speed | 24 | y = 20.18 (± 5.76) + 11.68 (± 2.27) x | 0.53 | < 0.001 |
| Iznájar 2018 | | | 24 | y = - 0.68 (± 1.53) + 2.37 (± 0.64) x | 0.36 | < 0.01 |
| Cubillas 2016 | | | 13 | not significant | | 0.617 |
| Cubillas 2018 | Diffusive $CH_4$ emissions | Water temperature | 24 | y = - 702.92 (± 219.97) + 28.95 (± 8.57) x | 0.31 | < 0.01 |
| Iznájar 2018 | | | 24 | y = - 57.08 (± 14.78) + 2.31 (± 0.56) x | 0.42 | < 0.001 |
| Cubillas 2016 | | Oxygen | 13 | not significant | | 0.478 |

 © xxxx IOP Publishing Ltd

| | | | | | | |
|---|---|---|---|---|---|---|
| Cubillas 2018 | Diffusive $CH_4$ emissions | saturation | 24 | y = - 674.19 (± 192.96) + 6.07 (± 1.64) x | 0.36 | < 0.01 |
| Iznájar 2018 | | | 24 | y = - 76.74 (± 21.87) + 0.61 (± 0.16) x | 0.36 | < 0.01 |


© xxxx IOP Publishing Ltd

Journal **XX** (XXXX) XXXXXX https://doi.org/XXXX/XXXX

**Table S7.** The fluxes of $CO_2$, $N_2O$, and $CH_4$ during the 24 h cycles in 2018, and during specific daytime periods. The mean and standard deviation (SD) are provided. The percentage between the mean for that time period, and the 24 h mean is provided in order to evaluate the over or underestimation of that specific time period in relation to the daily mean.

| | | $CO_2$, µmol m$^{-2}$ s$^{-1}$ (mean ± SD) | | $N_2O$, nmol m$^{-2}$ s$^{-1}$ (mean ± SD) | | Total $CH_4$, nmol m$^{-2}$ s$^{-1}$ (mean ± SD) | | Diffusive $CH_4$, nmol m$^{-2}$ s$^{-1}$ (mean ± SD) | | Ebullitive $CH_4$, nmol m$^{-2}$ s$^{-1}$ (mean ± SD) | |
|---|---|---|---|---|---|---|---|---|---|---|---|
| Cubillas reservoir, 2018 | 24 h cycle | 0.47 ± 0.35 | | 0.30 ± 0.29 | | 100.82 ± 52.81 | | 40.46 ± 29.80 | | 60.36 ± 44.03 | |
| | 9 – 5 pm | 0.44 ± 0.11 | 92 % | 0.24 ± 0.09 | 79 % | 86.23 ± 34.80 | 86 % | 34.63 ± 9.33 | 86 % | 51.60 ± 29.14 | 85 % |
| | 9 - 2 pm | **0.47 ± 0.12** | **98 %** | 0.28 ± 0.10 | 91 % | 90.85 ± 38.88 | 90 % | 37.44 ± 5.01 | 93 % | 53.41 ± 35.10 | 88 % |
| | 10 - 3 pm | 0.51 ± 0.12 | 107 % | **0.28 ± 0.09** | **93 %** | 95.70 ± 30.30 | 95 % | 38.39 ± 2.95 | 95 % | 57.31 ± 28.81 | 95 % |
| | 11 - 4 pm | 0.54 ± 0.09 | 115 % | 0.28 ± 0.08 | 92 % | 110.50 ± 36.38 | 110 % | **41.54 ± 10.42** | **103 %** | 68.96 ± 30.13 | 114 % |
| | 12 - 5 pm | 0.61 ± 0.17 | 129 % | 0.35 ± 0.15 | 115 % | **104.26 ± 26.93** | **103 %** | 44.42 ± 13.49 | 110 % | **59.84 ± 16.33** | **99 %** |
| Iznájar reservoir, 2018 | 24 h cycle | 0.17 ± 0.12 | | 0.17 ± 0.16 | | 9.45 ± 10.53 | | 4.43 ± 4.15 | | 5.02 ± 7.86 | |
| | 9 - 5 pm | 0.23 ± 0.08 | 135 % | 0.23 ± 0.13 | 132 % | 12.23 ± 10.73 | 129 % | 5.89 ± 4.10 | 133 % | 6.34 ± 8.10 | 126 % |
| | 9 - 2 pm | **0.17 ± 0.04** | **102 %** | 0.15 ± 0.05 | 86 % | 10.00 ± 10.62 | 106 % | 3.56 ± 1.55 | 80 % | 6.43 ± 9.19 | 128 % |
| | 10 - 3 pm | 0.20 ± 0.09 | 120 % | **0.20 ± 0.15** | **118 %** | **9.29 ± 10.32** | **98 %** | **4.14 ± 2.46** | **93 %** | 5.15 ± 9.01 | 103 % |
| | 11 - 4 pm | 0.23 ± 0.10 | 138 % | 0.24 ± 0.15 | 140 % | 9.99 ± 9.94 | 106 % | 4.95 ± 2.58 | 112 % | **5.04 ± 9.09** | **100 %** |
| | 12 - 5 pm | 0.27 ± 0.08 | 158 % | 0.29 ± 0.12 | 166 % | 15.55 ± 11.78 | 165 % | 7.52 ± 4.37 | 170 % | 8.03 ± 9.47 | 160 % |


© xxxx IOP Publishing Ltd

**Table S8.** Summary table of published studies on the variability of $CO_2$ (μmol m$^{-2}$ s$^{-1}$), $N_2O$ (nmol m$^{-2}$ s$^{-1}$), and $CH_4$ (nmol m$^{-2}$ s$^{-1}$) fluxes at daily scales in different ecosystems and using different methods. We classified these methods in direct measurement, by eddy covariance technique (EC); closed/floating chamber (FC) and bubble traps for ebullitive fluxes (BT); and the indirect method that predicted the fluxes (PF) based on gas transfer velocities (k). The transfer velocities are traditionally based on wind-speed, as $k_{CC}$ [13] and $k_{CW}$ for low wind speeds [14]. Other gas transfer velocities also include the effects of water-side cooling to the gas transfer besides shear-induced turbulence, as $k_{HE}$ [75] and $k_{TE}$ [76]. Other authors preferred an empirical model for k, derived from laboratory experiments. We provided the daytime and nighttime mean (value ± standard error); or the range of variation (minimum - maximum). We provided the daily values when the data for daytime/nighttime are not supplied in the reference study.

| GHG | Ecosystem, mean depth, and state | Method | Daytime flux | Nighttime flux | Study |
|---|---|---|---|---|---|
| $CO_2$ | Reservoir (70 - 130 m) | Direct: FC | August: 0.74 - 2.71<br>October: 0.27 - 1.46<br>February: 0.12 - 0.53<br>April: 0.14 - 1.74 | | [3] |
| | Reservoir (5 m, 1 year period) | Direct: EC | 0.23 | 0.39 | [4] |
| | Reservoir (littoral zone) | Direct: FC | 0.06 ± 0.04 | 0.13 ± 0.05 | [5] |
| | Lake (1.3 m, 2 years period) | Direct: EC | -0.78 - 1.62<br>Higher fluxes at nighttime | | [52] |
| | Lake (8 m, several seasons) | Direct: EC | 0.35 | 0.31 | [58] |
| | | Indirect: PF ($k_{TE}$ or $k_{CC}$) | $k_{TE}$: 0.31<br>$k_{CC}$: 0.28 | $k_{TE}$: 0.23<br>$k_{CC}$: 0.23 | |
| | Lake (6.3 m, stratified) | Direct: EC and FC | EC: 0.31 ± 0.04<br>FC: 0.62 ± 0.80 | EC: 0.28 ± 0.08<br>FC: 0.29 ± 0.04 | [59] |
| | | Indirect: PF | $k_{HE}$: 0.305 ± 0.009<br>$k_{TE}$: 0.545 ± 0.014<br>$k_{CC}$: 0.201 ± 0.004 | $k_{HE}$: 0.410 ± 0.008<br>$k_{TE}$: 0.396 ± 0.010<br>$k_{CC}$: 0.180 ± 0.004 | |
| | Lake (6.3 m, mixed) | Direct: EC | 1.3 ± 0.2 | 0.88 ± 0.14 | |
| | | Indirect: PF ($k_{HE}$, $k_{TE}$ or $k_{CC}$) | $k_{HE}$: 2.15 ± 0.06<br>$k_{TE}$: 2.37 ± 0.06<br>$k_{CC}$: 1.11 ± 0.04 | $k_{HE}$: 1.43 ± 0.05<br>$k_{TE}$: 1.54 ± 0.05<br>$k_{CC}$: 0.58 ± 0.02 | |
| | Lakes (stratified): Dagow (6 m, oligo-mesotrophic), Stechlin (21 m, eutrophic) | Direct: FC | Dw: 0.03 - 0.13<br>Higher daytime fluxes<br><br>St: - 0.10 - (- 0.04)<br>No differences found | | [41] |
| | Shallow lake (0.5 – 2.5 m) | Direct: FC | Higher fluxes at nighttime over seasons | | [77] |
| | Wetland (0.5 m) | Direct: EC | - 3.58 | 3.24 | [53] |
| | Cubillas 2016 (7.0 m), Cubillas 2018, (8.9 m), and Iznájar 2018 (22.4 m) | Direct: FC | C1: 0.12 - 0.42<br>C2: 0.26 - 1.46<br>Iz: 0.11 - 0.35 | C1: 0.10 - 0.21<br>C2: 0.17 - 0.39<br>Iz: -0.01 - 0.27 | This study |

 © xxxx IOP Publishing Ltd

| | | | | | |
|---|---|---|---|---|---|
| $N_2O$ | River<br>(0.01 - 0.15 m) | Indirect:<br>PF | Daily mean: 5.96<br>Mean at daytime : 9.69 | | [31] |
| | River<br>(1.2 m) | Direct: FC<br>Indirect: PF | FC: 0.52 - 1.39<br>PF: 0.13 - 0.25<br>Higher fluxes at daytime | | [32] |
| | Rivers<br>(FLR, HBR, NFR) | Direct: FC | FLR: 0.05 - 0.10<br>HBR: 0.04 - 0.11<br>NFR: 0.45 - 0.94 | FLR: 0.05 - 0.06<br>HBR: 0.08 - 0.09<br>NFR: 0.70 - 1.03 | [19] |
| | 10 Streams<br>(0.10 - 0.60 m) | Indirect:<br>PF | 0.94 | 0.86 | [30] |
| | River<br>(23 sites, < 1 m) | Indirect:<br>PF | Higher fluxes at nighttime, numbers not provided | | [78] |
| | River<br>(3 sites,<br>13 - 19 m) | Indirect:<br>PF | Daily mean S1: 0.08 - 0.25<br>Daily mean S2: 0.02 - 0.35<br>Daily mean S3: 0.08 - 0.40<br>Peak of emission at daytime | | [55] |
| | River<br>(2.5 m) | Direct: FC | 0.41 - 0.87<br>0.61 | 0.50 | [20] |
| | Grassland soil (0.15 m, different season) | Direct: FC | June: 0.01 - 0.11<br>July: 0.05 - 0.62<br>August: 0.03 - 0.11<br>September: 0.00 - 0.03<br>Peaks of emission at daytime | | [54] |
| | Soil<br>Mesocosms | Direct: FC | Treatments presented different daily means and ranges, but all treatments had higher fluxes at daytime | | [61] |
| | Cubillas 2016 (7.0 m),<br>Cubillas 2018, (8.9 m),<br>and Iznájar 2018 (22.4 m) | Direct: FC | C1: 0.03 - 0.31<br>C2: 0.13 - 1.13<br>Iz: 0.07 - 0.47 | C1: 0.03 - 0.11<br>C2: 0.08 - 0.24<br>Iz:-0.09 - 0.28 | This study |
| $CH_4$ | Reservoir<br>(1 - 10 m) | Direct:<br>FC and BT | FC: 523 ± 479<br>(4 - 1502)<br>BT: 849 ± 192 | FC: 336 ± 442<br>BT: 260 ± 77 | [17] |
| | Reservoir<br>(70 - 130 m) | Direct: FC | August: 0.5 - 2.1<br>October: 0.2 - 2.3<br>February: 0.0 - 0.5<br>April: 0.2 - 5.5 | | [3] |
| | Reservoir (12 m, eutrophic)<br>2 sites (S1, S2) | Direct: FC | Jan, S1: 1.0 ± 0.4<br>Jan, S2: 0.8 ± 0.2<br>Apr S1: 3.8 ± 2.4<br>Apr S2: 1.4 ± 0.9<br>July S1: 4.1 ± 1.9<br>July S2: 2.8 ± 1.5 | Jan, S1: 0.8 ± 0.4<br>Jan, S2: 0.7 ± 0.2<br>Apr S1: 1.4 ± 0.9<br>Apr S2: 1.1 ± 0.5<br>July S1: 4.2 ± 2.3<br>July S2: 1.2 ± 0.6 | [56] |
| | 73 Lakes | Direct: FC | 9 - 158 % greater emissions<br>during the day | | [62] |
| | 16 Lakes<br>(0.5 - 3.4 m) | Direct: FC | The daily means ranged from 45 - 859<br>Daytime fluxes were significantly higher | | [11] |
| | Lake<br>(1.3 m) | Direct: EC | 0 - 300<br>Highest values during night<br>and early morning | | [22] |


© xxxx IOP Publishing Ltd

| | | | | | |
|---|---|---|---|---|---|
| | Lake (6.3 m, stratified) | Direct: EC and FC | EC: 0.41 ± 0.04<br>FC: 2.4 ± 0.3 | EC: 0.34 ± 0.04<br>FC: 1.1 ± 0.2 | [59] |
| | | Indirect: PF ($k_{HE}$, $k_{TE}$ or $k_{CC}$) | $k_{HE}$: 0.177 ± 0.005<br>$k_{TE}$: 0.370 ± 0.011<br>$k_{CC}$: 0.128 ± 0.003 | $k_{HE}$: 0.431 ± 0.008<br>$k_{TE}$: 0.439 ± 0.007<br>$k_{CC}$: 0.186 ± 0.004 | |
| | Lake (6.3 m, mixed) | Direct: EC | 5.9 ± 0.3 | 5.0 ± 0.4 | |
| | | Indirect: PF ($k_{HE}$, $k_{TE}$ or $k_{CC}$) | $k_{HE}$: 7.1± 0.6<br>$k_{TE}$: 7.7 ± 0.6<br>$k_{CC}$: 3.7 ± 0.3 | $k_{HE}$: 6.6 ± 0.5<br>$k_{TE}$: 7.1 ± 0.5<br>$k_{CC}$: 2.8 ± 0.2 | |
| | Lakes (stratified): Dagow (6 m, oligo-mesotrophic), Stechlin (21 m, eutrophic) | Direct: FC | D: 7.7 - 49.3<br>S: 0.7 - 1.5<br>No differences found between daytime and nighttime fluxes | | [41] |
| | 4 lakes (<0.5 – 8 m) | Direct: FC | Higher emissions at daytime than at nighttime | | [8] |
| | Shallow lake (0.5 – 2.5 m), Two sites (S1, S2) over several months | Direct: FC | Higher fluxes at nighttime over seasons<br>S1, diffusive: 103.9 (1.1 – 450.0)<br>S2, diffusive: 119.7 (0.8 – 697.5) | | [57] |
| | Wetland: bog (0.2 m) | Direct: EC | 15.0 ± 18.7 | 6.9 ± 6.2 | [16] |
| | Wetland (0.16 - 0.32 m) | Indirect: PF | 126 ± 21 | 75 ± 11 | [63] |
| | Wetland (0.5 m) | Direct: EC | 30 - 1600<br>Peak of emission at daytime | | [53] |
| | Pond (1.45 m) | Direct: FC | Diffusive: 3.1<br>Ebullitive: 8.1 | Diffusive: 3.1<br>Ebullitive: 5.0 | [18] |
| | Cubillas 2016 (7.0 m), Cubillas 2018, (8.9 m), and Iznájar 2018 (22.4 m) | Direct: FC | Diffusive fluxes<br>C1: 0 - 989<br>C2: 17 - 143<br>Iz: 2 - 15<br>Ebullitive fluxes<br>C1: 0 - 1815<br>C2: 1 - 207<br>Iz: 0 - 21 | Diffusive fluxes<br>C1: 27 - 169<br>C2: 11 - 36<br>Iz: 0 - 9<br>Ebullitive fluxes<br>C1: 133 - 920<br>C2: 16 - 97<br>Iz: 0 - 27 | This study |

70. Beaulieu, J.J.; Balz, D.A.; Birchfield, M.K.; Harrison, J.A.; Nietch, C.T.; Platz, M.C.; Squier, W.C.; Waldo, S.; Walker, J.T.; White, K.M.; et al. Effects of an Experimental Water-Level Drawdown on

 © xxxx IOP Publishing Ltd

Methane Emissions from a Eutrophic Reservoir. *Ecosystems* **2018**, *21*, 657–674, doi:10.1007/s10021-017-0176-2.

 © xxxx IOP Publishing Ltd
